# Nanoparticle-Conjugated TLR9 Agonists Improve the Potency, Durability, and Breadth of COVID-19 Vaccines

**DOI:** 10.1101/2023.01.02.522505

**Authors:** Ben S. Ou, Julie Baillet, Vittoria C.T.M. Picece, Emily C. Gale, Abigail E. Powell, Olivia M. Saouaf, Jerry Yan, Anahita Nejatfard, Hector Lopez Hernandez, Eric A. Appel

## Abstract

Development of effective vaccines for infectious diseases has been one of the most successful global health interventions in history. Though, while ideal subunit vaccines strongly rely on antigen and adjuvant(s) selection, the mode and timescale of exposure to the immune system has often been overlooked. Unfortunately, poor control over the delivery of many adjuvants, which play a key role in enhancing the quality and potency of immune responses, can limit their efficacy and cause off-target toxicities. There is critical need for new adjuvant delivery technologies to enhance their efficacy and boost vaccine performance. Nanoparticles have been shown to be ideal carriers for improving antigen delivery due to their shape and size, which mimic viral structures, but have been generally less explored for adjuvant delivery. Here, we describe the design of self-assembled poly(ethylene glycol)-*b-*poly(lactic acid) nanoparticles decorated with CpG, a potent TLR9 agonist, to increase adjuvanticity in COVID-19 vaccines. By controlling the surface density of CpG, we show that intermediate valency is a key factor for TLR9 activation of immune cells. When delivered with the SARS-CoV-2 spike protein, CpG nanoparticle (CpG-NP) adjuvant greatly improve the magnitude and duration of antibody responses when compared to soluble CpG, and result in overall greater breadth of immunity against variants of concern. Moreover, encapsulation of CpG-NP into injectable polymeric-nanoparticle (PNP) hydrogels enhance the spatiotemporal control over co-delivery of CpG-NP adjuvant and spike protein antigen such that a single immunization of hydrogel-based vaccines generates comparable humoral responses as a typical prime-boost regimen of soluble vaccines. These delivery technologies can potentially reduce the costs and burden of clinical vaccination, both of which are key elements in fighting a pandemic.

## INTRODUCTION

Vaccines are among the most effective medical advancements in history and are estimated to save 2.5 million lives worldwide annually.^1^ Unfortunately, an abundance of infectious diseases – including many rapidly mutating viral pathogens such as HIV, influenza, and SARS-CoV-2 – still do not have sufficiently effective vaccines capable of providing broad and durable protection for a global population. To date, roughly one million people die each year from flu and HIV and COVID has killed more than 6.5 million people since its arrival three years ago^2^, highlighting the continuing threat of pandemic viruses and a critical need for improved vaccine technologies.

Among the common types of vaccines used in the clinic, subunit vaccines offer excellent safety, stability, scalability, and worldwide manufacturing capabilities, as well as more widely available storage conditions compared to mRNA-based vaccines.^3^ Subunit vaccines contain protein antigens, which direct the antibody response to a specific foreign substance, along with one or more immune stimulating additives commonly referred to as adjuvants. These adjuvant materials have been shown to play a key role in enhancing the body’s immune response to a pathogen and therefore vaccine efficacy. Yet, there is a growing need for new approaches to augment adjuvant potency and enhance the quality and durability of immune responses.^4^ Some of the most widely used clinical adjuvants include aluminum salt-based adjuvants (Alum) and squalene-based oil-in-water emulsions such as MF59 and AS03, though the specific mechanism of action of these adjuvants is poorly understood.^5^ There are also several molecular adjuvants which trigger innate immune cell activation through signaling of pattern recognition receptors (PRR), including toll-like receptor agonists (TLRas) such as oligodeoxynucleotide CpG ODN (TLR9 agonist) and MPL (TLR4 agonist). CpG, for example, has been shown to increase immune responses by strongly activating innate immune cells as the CpG motifs mimic the activity of bacterial DNA.^6^ Specifically, CpG triggers intracellular signaling leading to the activation of antigen presenting cells (APCs) such as macrophages and dendritic cells (DCs) and B cells, as well as the production of chemokines and cytokines, enhancing both innate and adaptive immune responses.^7–14^ CpG is included in the Hepatitis B vaccine Heplisav B (FDA approved in 2017), and is also currently being evaluated clinically in multiple SARS-CoV-2 vaccines.^15^

To maximize the activation of innate immune cells while minimizing systemic toxicities,^16, 17^ significant efforts have focused on localizing TLR agonists to the injection site and lymph nodes (LNs). Nanoparticles (NPs) have been extensively explored as delivery carriers due to their modularity, scalability, biocompatibility, and ability to overcome spatiotemporal challenges associated with conventional delivery methods.^16–29^ NPs between 20 and 100 nm^30–35^ efficiently drain through the lymphatic system into the LNs^30, 31, 36^ where they are directly taken up by LN-resident APCs^37^ without requiring specific cell targeting ligands. Moreover, recent studies have reported that covalently conjugating TLRa molecules to polymer NPs resulted in a significant increase in both antibody production and induction of cytotoxic T-cells.^17^ Similarly, other studies have leveraged particle technologies such as liposomes and lipid nanoparticles for delivery of multiple adjuvant molecules, including CpG, to improve the potency of the adjuvant response.^13, 17, 38–45^ Yet, these technologies have exhibited limitations in either the complexity of manufacturing, challenges with scalability, poor control over TLRa valency or dosing, or limited range of available particle sizes. Indeed, while liposomes and lipid nanoparticles can be manufactured in sizes typically ranging from 80-150nm, it has been reported that smaller particles are better taken up by important APCs such as DCs.^37^ ^46^ Furthermore, a growing body of work demonstrated that immune cells require precise spatial and temporal cues to drive specified responses, supporting the idea that spatiotemporal control of vaccines can have a profound effect on the magnitude and quality of the immune response.^47–56^

The precise spatiotemporal control of vaccines can also be achieved by prolonged, localized co-delivery of vaccine components. Recent studies have shown that sustained release of an HIV vaccine through implantable osmotic pumps led to greatly improved quality of vaccine responses, such as durable germinal center (GC) responses, high antibody titers, and development of better virus neutralization compared to standard soluble administration of the same vaccine.^53^ Similarly, microneedles and injectable hydrogels have been widely used as slow vaccine delivery platforms.^29^ Our group has developed injectable polymer-nanoparticle (PNP) hydrogels for prolonged co-delivery of subunit vaccine components.^50, 57–61^ We have demonstrated that PNP hydrogels can provide sustained release of distinct vaccine cargo over the course of weeks, while prolonging the GC reaction and improving antibody affinity by more than 1000-fold compared to soluble vaccine formulation.^50, 62, 63^

In the current study, we sought to optimize the delivery of CpG to improve the potency by developing a nanoparticle-based adjuvant construct. We chemically conjugated the CpG adjuvant to poly(ethylene glycol)-*b-*poly(lactic acid) (PEG-*b*-PLA) NPs (CpG-NPs) of approximately 50 nm in size, allowing for efficient, passive and direct transport to the LNs (Figure 1). We showed that the bioactivity of CpG was not affected after conjugation and that precise tuning of the CpG valency on the NPs surface enabled control of the potency of the elicited immune response *in vitro*. Moreover, these modifiable CpG-NPs can be embedded into PNP hydrogels for sustained exposure of vaccine adjuvants to improve immune responses. In this regard, we compared the adjuvanticity of soluble CpG, CpG-NPs, and PNP hydrogels containing CpG-NPs (CpG-NP hydrogels) *in vivo* as part of a COVID-19 subunit vaccine using the SARS-CoV-2 spike protein as antigen. We showed that a single immunization of CpG-NP hydrogel as well as a prime-boost soluble CpG-NP vaccine demonstrated superior anti-spike antibody titers and broader antibody responses against immune-evading variants. Overall, we report the facile design of a broadly implementable CpG-NP platform that can improve adjuvant potency leading to increased breadth and durability of vaccines.

**Figure 1:**
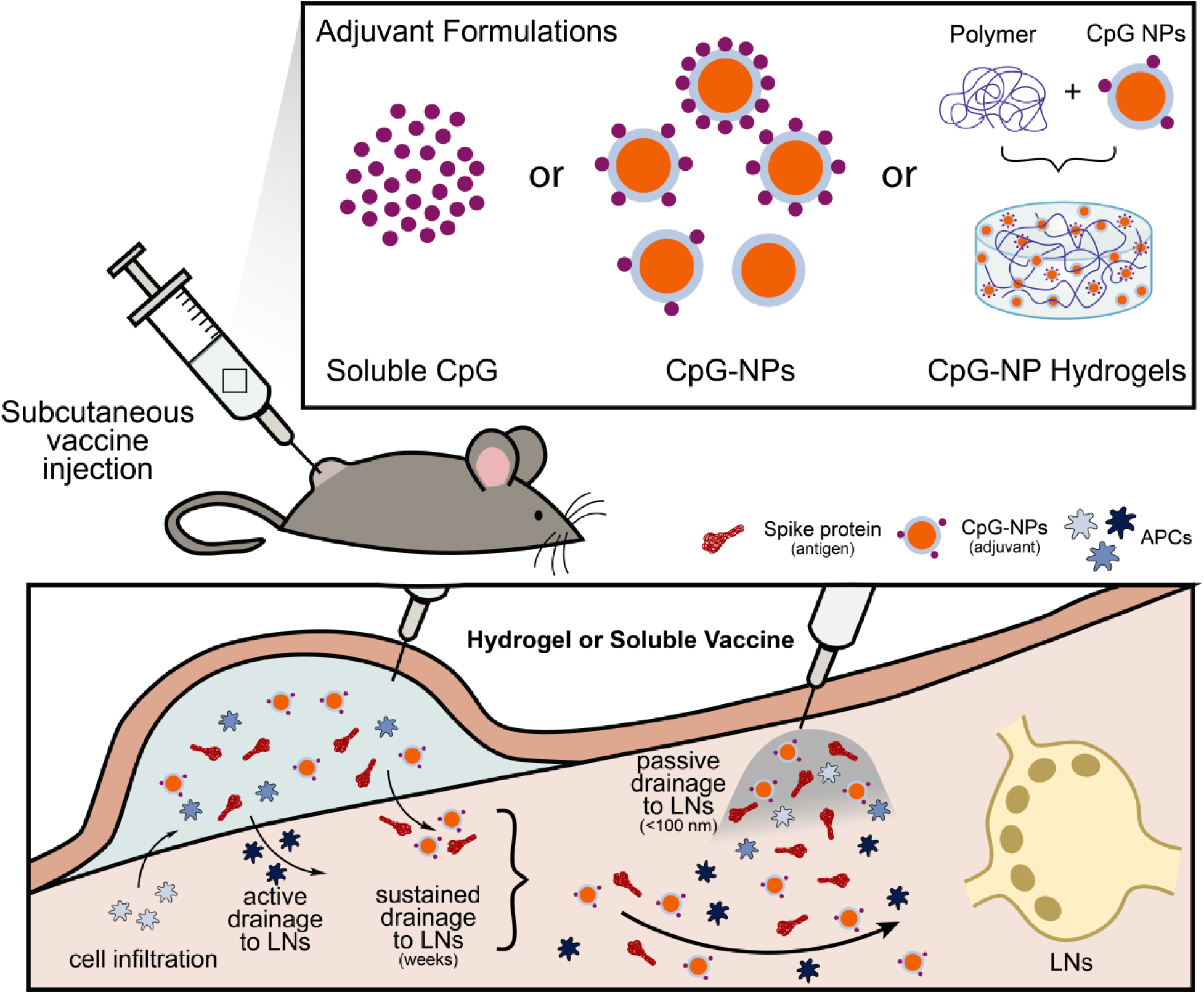
Schematic representation of a subcutaneous vaccine injection in a mouse model for in vivo release. Delivery of CpG adjuvant can be achieved in different ways: in its molecular form, tethered to PEG-*b*-PLA NPs, or tethered to NPs and encapsulated in polymer-nanoparticle (PNP) hydrogels. PNP hydrogels are loaded with vaccine cargo, including antigen and adjuvant (CpG-NPs), and allow for sustained vaccine exposure. After subcutaneous injection of the hydrogel vaccine, vaccine components can be transported to the lymph nodes (LNs) either by drainage through antigen presenting cells (APCs) that have previously infiltrated the hydrogel, or by LN drainage of the single vaccine components themselves. Soluble vaccines, on the other hand, do not create an inflammatory niche for cell infiltration. Vaccine components are rapidly cleared from the body and drained to the lymph nodes, potentially decreasing the potency. Nanoparticle vaccine cargo, such as CpG-NPs, however, may improve immune cells activation and LN targeting ability.

## RESULTS AND DISCUSSION

### Synthesis and characterization of CpG-NPs

We conjugated the TLR9 adjuvant CpG to PEG-*b*-PLA NPs to improve its stability, its targeting to LNs and its uptake by APCs (Figure 2). We have previously described the synthesis of azide-terminated PEG-*b*-PLA (N_3_-PEG-*b*-PLA) block copolymers using organocatalytic ring-opening polymerization (ROP) and their self-assembly in core-shell types NPs.^26, 27, 57, 58, 64, 65^ DBCO-modified CpG was tethered on the surface of N_3_-PEG-*b*-PLA NPs using copper-free strain-promoted cycloaddition (Figure 2A). Conversions higher than 90% were obtained with a threefold molar excess of DBCO-CpG with respect to the azide functionality on the NPs (Figure S1). This modular approach allows the use of various classes and sequences of CpG. In this work, the CpG-2395 sequence, belonging to class C CpG (CpG-C, abbreviated to CpG), was selected due to its ability to activate both human and murine immune cells, thereby reinforcing translational efforts towards potential pre-clinical applications. Furthermore, CpG-C were found to both strongly induce plasmacytoid DCs (pDCs) in secreting IFN-α and TNF-α as well as B cells activation and proliferation.^66–71^

**Figure 2:**
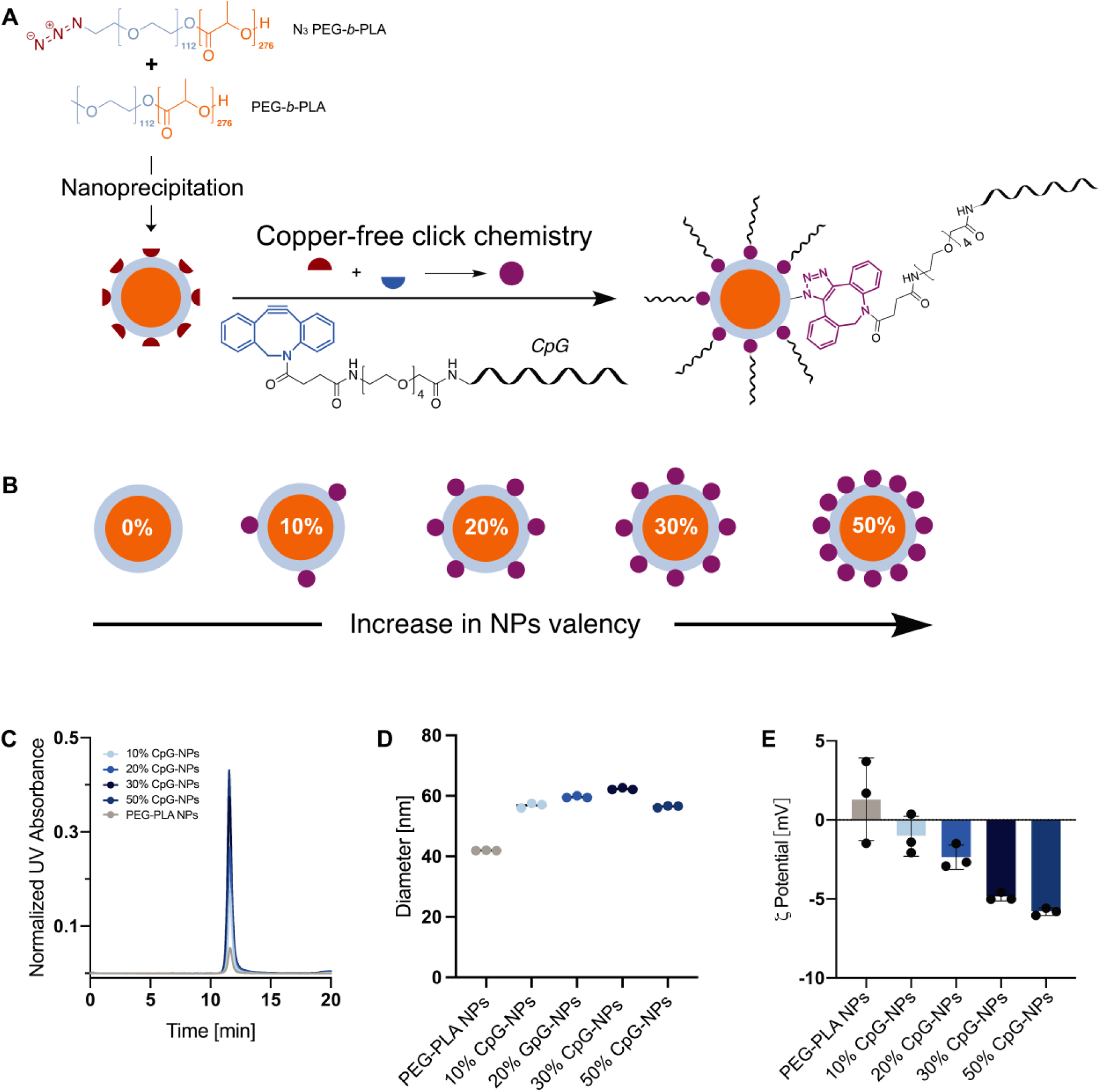
Design of CpG functionalized NPs. **(A)** Synthetic scheme for the fabrication of CpG-based NPs. Formation of azide-terminated PEG-*b*-PLA NPs via nanoprecipitation followed by copper-free click chemistry with DBCO-CpG to yield to CpG-functionalized NPs. 10%, 20%, 30% and 50% valencies were achieved by mixing different weight ratios of PEG-*b*-PLA and N_3_-PEG-*b*-PLA polymer solutions before nanoprecipitation. **(B)** Normalized UV absorbance of 10%, 20%, 30% and 50% CpG-functionalized NPs. **(C)** Hydrodynamic diameters of PEG-*b*-PLA NPs and CpG-NPs in PBS 1X. **(D)** Surface zeta potential of PEG-*b*-PLA NPs and CpG-NPs in PBS 1X.

Different valencies of TLR agonists on NPs have been shown to influence the magnitude and persistence of innate immune activation by leading to higher expression of co-stimulatory molecules.^17, 65^ Therefore, manipulating the density of the CpG adjuvant molecules on the surface of the PEG-*b*-PLA NPs could potentially improve the potency of adjuvant responses.^29, 38, 65^ A series of NPs with increasing CpG valencies on the surface were obtained by controlling the number of azide functionalities when physically mixing different weight ratios of N_3_-PEG-*b*-PLA and PEG-*b*-PLA (i.e., 10%, 20%, 30% and 50%; SI). The resulting CpG-NPs were purified via size exclusion chromatography and the purity was assessed by size exclusion chromatography and gel electrophoresis (Figures S2, S3). The conjugation did not affect the physical and colloidal properties of the NPs and successful CpG functionalization was confirmed by an increase in UV absorbance, hydrodynamic diameter, and the zeta potential (Figures 2B, 2C, 2D). CpG-NPs were found to have hydrodynamic diameters between 56 to 62 nm (Table S1), which is within the size range known to demonstrate improved trafficking to LNs^30,31,36^ while avoiding immediate partitioning of soluble CpG into the blood stream and therefore reducing systemic toxicities (Figures 2C, S4). Moreover, the negatively charged phosphorothioate backbone of CpG induced an increase of negative charge on the NPs, therefore increasing the colloidal stability of the NPs (Figure 2D).

### In vitro and in vivo evaluation of CpG-NPs’ cellular activation, uptake, and biodistribution

To ensure that CpG conjugation to the NPs did not impair the biological activity and immunogenicity of the adjuvant, RAW-Blue transgenic mouse macrophage cells and human THP-1 hTLR9 monocyte cells were used to quantify TLR9 activation (Figure 3a). The cells were incubated with either CpG-NPs or soluble CpG. Additionally, this *in vitro* assay was used to evaluate the effect of CpG valency on the potency of innate immune cell activation. In these assays, Raw-Blue cells were incubated for 21 h with either soluble CpG, plain PEG-*b-*PLA NPs, or CpG-conjugated NPs (with valencies of 10%, 20%, 30%, 50%) at a range of CpG concentrations (3.1– 29 μg/mL) to generate concentration-dependent activation curves (Figure 3B). To ensure a correct and consistent dosing of CpG-NPs throughout different experiments, we constructed a standard curve for the CpG-NPs (Figure S5). From the normalized dilution curves, we then determined the EC_50_ values (Figures 3C, S6). We observed that CpG density influenced the resulting EC_50_ values and therefore the overall potency. 30% CpG-NPs resulted in the lowest EC_50_ value (log EC_50_ = 1.3 μg/mL) compared to all other valencies and behaved similarly to soluble CpG (log EC_50_ = 1.18 μg/mL). Interestingly, 50% valency resulted in the highest EC_50_ value (log EC_50_ = 2.0 μg/mL), suggesting a low level of TLR9 activation. The decrease in potency observed with the highest CpG valency (50% CpG-NPs) could be due to CpG saturation and a highly negatively charged surface which, by surpassing the critical threshold for charge density, could decrease CpG accessibility. We further verified this finding with a second *in vitro* activation assay using human THP-1 hTLR9 monocyte cells (Figure 3A). THP-1 cells were incubated with either 30% CpG-NPs, 50% CpG-NPs, or soluble CpG at a range of CpG concentrations (3.1-29 μg/mL) to generate concentration curves (Figure 3D). 30% CpG-NPs resulted in the highest activation signal at a CpG concentration of 29 μg/mL, thereby supporting their higher potency over 50% CpG-NPs (Figure 3E). These data demonstrate that we were able to synthesize CpG-NPs of different valencies, with the intermediate CpG valency of 30% activating TLR9 similarly to soluble CpG *in vitro*.

**Figure 3:**
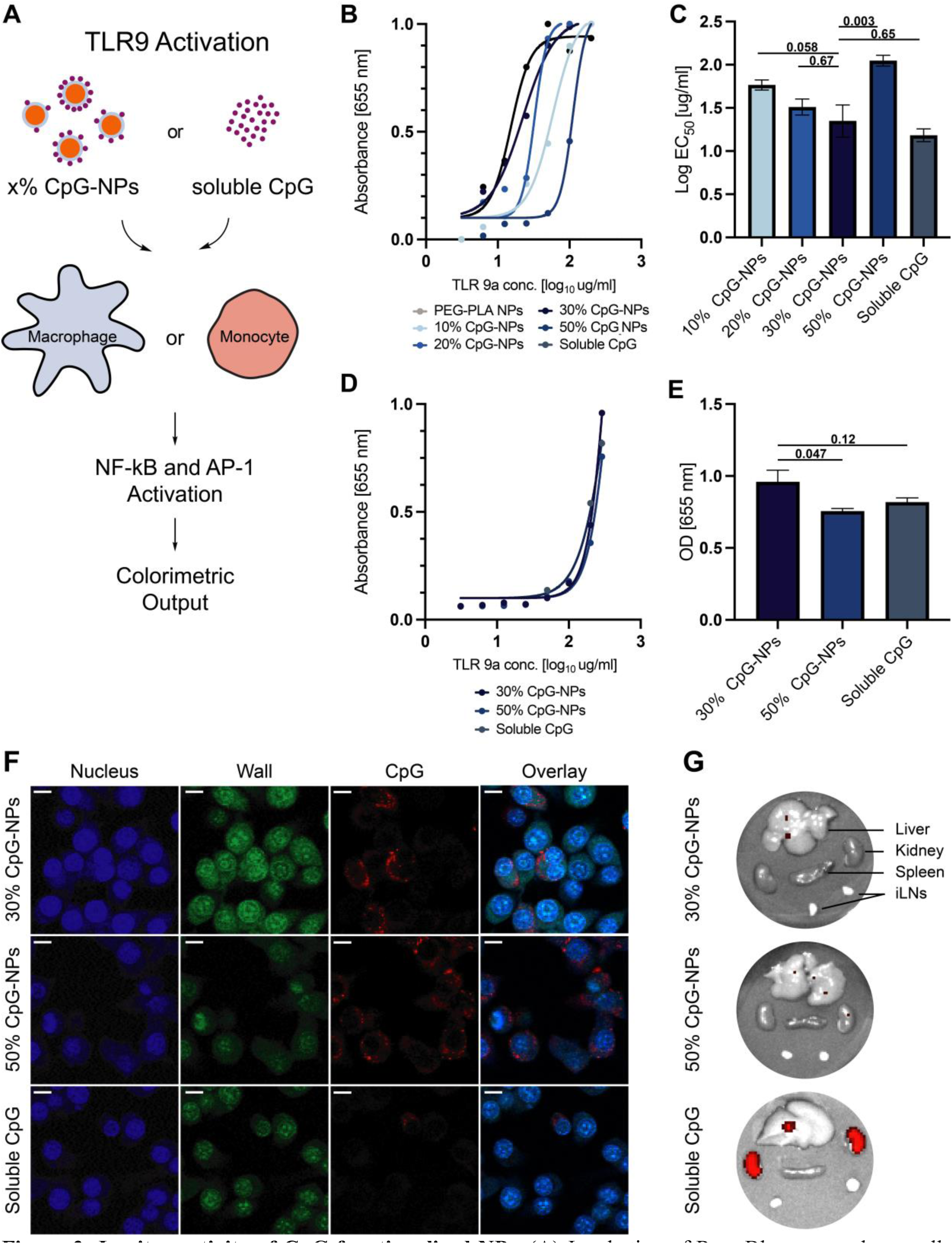
In vitro activity of CpG functionalized NPs. **(A)** Incubation of Raw-Blue macrophage cells and THP1 hTLR9 monocyte cells with either soluble CpG or different valencies of CpG-NPs (10%, 20%, 30%, 50%) induces the activation of NF-kB and AP-1. The magnitude of activation is quantified via colorimetric output using QUANTI-Blue solution. **(B)** Normalized activation curves across a range of CpG concentrations (3.1-29 µg/mL) delivered on CpG-NPs at different densities to 100,000 Raw-Blue cells. The absorbance at 655 nm corresponds to TLR activation. **(C)** Log EC_50_ values for each activation curve were extrapolated from **(B)** using a “log(TLR9 agonist) vs response” nonlinear regression curve fit of the dilution curves. **(D)** Activation curves across a range of CpG concentration (3.1-29 µg/mL) delivered with different CpG formulations to 100,000 THP1-dual hTLR9 cells. **(E)** Optical density of different CpG formulations at a CpG concentration of 29 μg/mL at 655 nm. **(F)** Confocal microscopy images of cellular uptake of Raw-Blue cells incubated with different CpG formulations equivalent to 5 μg of CpG. Cell nucleus was stained with DAPI, cell wall was stained with Alexa Fluor 488 Anti-alpha 1 Sodium Potassium ATPase antibody, and CpG was conjugated with Cy5. Scale bars are 10 μm. **(G)** Accumulation of Cy5-conjugated CpG in organs of interest 3 h after injection. Images and signal were determined by an In Vivo Imaging System. *p* values listed were determined using a 1way ANOVA with Tukey’s multiple comparisons test. *p* values for comparisons between the 30% CpG-NPs group and all other groups are shown above the bars.

To assess cellular uptake and biodistribution of CpG-NPs, we synthesized fluorescently tagged CpG-NPs by tethering DBCO-modified Cy5-CpG on the surface of N_3_-PEG-*b*-PLA NPs using a similar synthetic route. We then incubated 5 μg CpG equivalent of 30% Cy5-CpG-NPs, 50% Cy5-CpG-NPs, or soluble Cy5-CpG with RAW-Blue macrophages as a model APC overnight to assess their uptakes. Confocal microscopy imaging confirmed that both CpG-NPs resulted in higher CpG internalization and co-localization than soluble CpG by counterstaining cells with DAPI and a surface stain (Figure 3F).

We then assessed the biodistribution of CpG-NPs compared to soluble CpG by subcutaneously injecting C57BL/6 mice with saline solutions of either 30% Cy5-CpG-NPs, 50% Cy5-CpG-NPs, or soluble Cy5-CpG (20 μg of CpG total for all formulations). We euthanized mice 3h post-injection and harvested their major organs (liver, kidney, spleen, and ipsilateral lymph nodes) to measure the distribution of CpG using an In Vivo Imaging System (IVIS). Mice injected with soluble CpG exhibited high accumulation of CpG in the kidney, consistent with previous findings reporting rapid systemic clearance of molecules below 20 kDa.^72, 73^ On the contrary, very little CpG accumulation was observed in the organs of animals dosed with either 30% CpG-NPs or 50% CpG-NPs, suggesting that CpG-NPs are better retained at the injection site and drained to the LNs. Based on both the *in vitro* and *in vivo* assessments of cellular activation, uptake, and distribution, 30% CpG-NPs was identified as the most promising adjuvant compared to other CpG valencies and soluble CpG to promote high magnitude of immune activation and persistent uptake and retention. We therefore selected 30% CpG-NPs as an adjuvant in a SARS-CoV-2 vaccine study, denoted simply as CpG-NPs in the following sections.

### Formulations of CpG-NP hydrogels and rheological characterization

Recent studies have highlighted the importance of sustained delivery in vaccines to prolong GC responses leading to improved breadth and affinity of antibody responses.^53, 74^ We have previously described the development of tunable and injectable PNP hydrogels able to encapsulate physiochemically diverse vaccine components such as antigens and adjuvants and to provide sustained co-delivery over extended periods of time.^50, 62, 63, 75^ We hypothesized that these unique characteristics could be coupled with the newly designed CpG-NPs featuring improved potency and *in vivo* trafficking properties to further enhance the vaccine response. PNP hydrogels can be easily formed by mixing aqueous solutions of hydrophobically modified hydroxypropyl methylcellulose derivatives (HPMC-C_12_) and biodegradable PEG-*b*-PLA NPs (Figure 4A).^50, 57, 60, 61^ After mixing, dynamic and multivalent non-covalent interactions form between the HPMC-C_12_ and the PEG-*b*-PLA NPs yielding dynamic crosslinks that lead to the formation of robust physical hydrogels (Figure 4B). These supramolecular hydrogels exhibit solid-like properties under static conditions, liquid-like behaviors under high shear, and rapid self-healing after succession of shear, allowing them to be readily injected through standard needles and form solid depots after injection.

**Figure 4:**
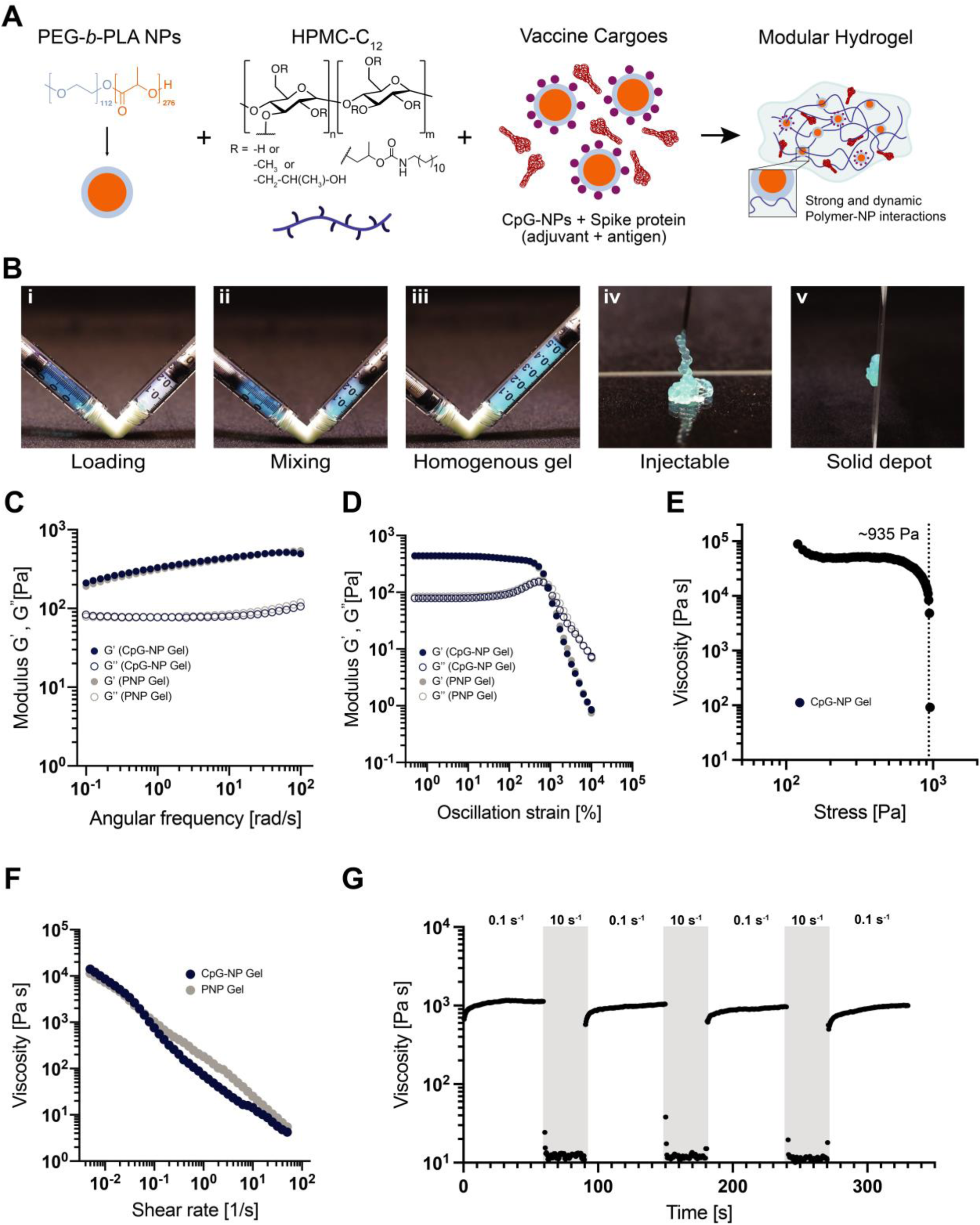
Fabrication and characterization of CpG-Polymer-Nanoparticle hydrogels. **(A)** Vaccine loaded CpG-NP hydrogels are formed when aqueous solutions of PEG-*b*-PLA NPs and dodecyl-modified hydroxypropylmethylcellulose (HPMC-C_12_) are mixed together with aqueous solutions of vaccine cargo comprising CpG-NPs (adjuvant) and spike protein (antigen). **(B)** Vaccine cargoes are added to the aqueous NPs solution before loading the aqueous and polymer components in two separate syringes (i), mixing the two phases with an elbow mixer (ii) leads to homogeneous hydrogels (iii). Image of a PNP hydrogel flowing through a 21-gauge needle during injection (iv) and formation of solid-like depot after injection (v). **(C)** Frequency-dependent oscillatory shear rheology and oscillatory amplitude sweeps **(D)** of CpG-NP and unloaded PNP hydrogels. **(E)** Stress-controlled flow sweeps of the CpG-NP hydrogel and yield stress value. **(F)** Shear-dependent viscosities of the two analyzed hydrogels demonstrate shear thinning and yielding properties, decreasing with increased shear rate. **(G)** Step-shear measurements over 3 cycles model yielding and healing of the hydrogels. Alternating low shear rates (0.1 1/s), and high shear rates (10.0 1/s, gray color) are imposed for 60 and 30 s respectively.

Vaccine components, including both adjuvants and antigens, can easily be loaded within the hydrogel network by simply mixing into the aqueous stock solutions during hydrogel manfacturing.^50^ To ensure that CpG-conjugation to the PEG-*b*-PLA NPs did not influence the mechanical properties of PNP hydrogels, we compared rheological properties of PNP hydrogels comprising CpG-NPs with standard hydrogel formulations (Figures 4C-G). We specifically investigated PNP hydrogel formulations comprising 2wt% HPMC-C_12_ and 10wt% NPs (containing a mixture of plain PEG-*b*-PLA NPs and CpG-NPs), which is denoted PNP-2-10.

Frequency-dependent oscillatory shear experiments were conducted within the linear viscoelastic regime (LVER) of the materials to measure their viscoelastic response. These experiments indicated that the introduction of CpG-NPs did not significantly alter the PNP hydrogel’s mechanical properties. Both formulations with and without CpG-NPs showed solid-like properties within the explored frequency range in which the storage (G’) modulus was greater than the loss (G’’) modulus (Figure 4C-4D). We also evaluated the yielding response of the hydrogels, which is an important characteristic for injectability and depot formation,^76^ using amplitude-dependent oscillatory shear experiments and stress-controlled flow experiments. Yield stress values of ∼935 Pa were measured by stress-controlled flow sweeps (Figure 4E). Further, the flow sweeps demonstrated that these materials exhibit a high degree of shear-thinning, whereby the measured viscosities decreased several orders of magnitude with increasing shear (Figure 4F).

Step-shear experiments were also conducted by interchanging in a stepwise fashion between low (0.1 1/s) and high (10 1/s) shear rates to determine self-healing behaviors of the hydrogels. The viscosity was observed to decrease by several orders of magnitude upon application of high shear rates, and rapidly and completely recovered when subjected to low shear rates (Figure 4G). From these observations, we confirmed that the inclusion of CpG-NPs did not alter the rheological characteristics of PNP hydrogels. We also showed these CpG-NP comprising hydrogels can be readily injected through high-gauge needles (Figure 4Biv) and maintain a robust structure after injection to allow for the formation of a robust depot *in vivo*.^50, 77^

### Vaccine cargo dynamics in PNP hydrogels

Vaccine components, including both antigens and adjuvants, typically exhibit highly distinct physiochemical properties that pose a challenge for their controlled and sustained delivery. These components can have extremely different polarities, charges, molecular weights, and hydrodynamic radii (*R_H_*) that may impact their encapsulation and diffusivity within a hydrogel network.^50^ Given the hydrophilicity and smaller molecular size of soluble CpG compared to the mesh size of a typical PNP-2-10 hydrogel formulation, soluble CpG has been previously shown to rapidly diffuse out of the matrix (half-life of release ∼2.5 days).^63, 78, 79^ We therefore hypothesized that sustained co-delivery of CpG-NPs and the SARS-CoV-2 spike protein, whose hydrodynamic sizes are expected to be much larger than the mesh size of the hydrogel, can be achieved with these materials. To evaluate this hypothesis, PNP-2-10 hydrogels were prepared with CpG-NPs (*R_H_ ∼* 30 nm) and loaded with spike protein (*R_H_ =* 12 nm, M*_w_* = 139 kDa). To characterize the dynamics of vaccine diffusion within the PNP hydrogels, we performed fluorescence recovery after photobleaching experiments (FRAP) (Figure 5).

**Figure 5:**
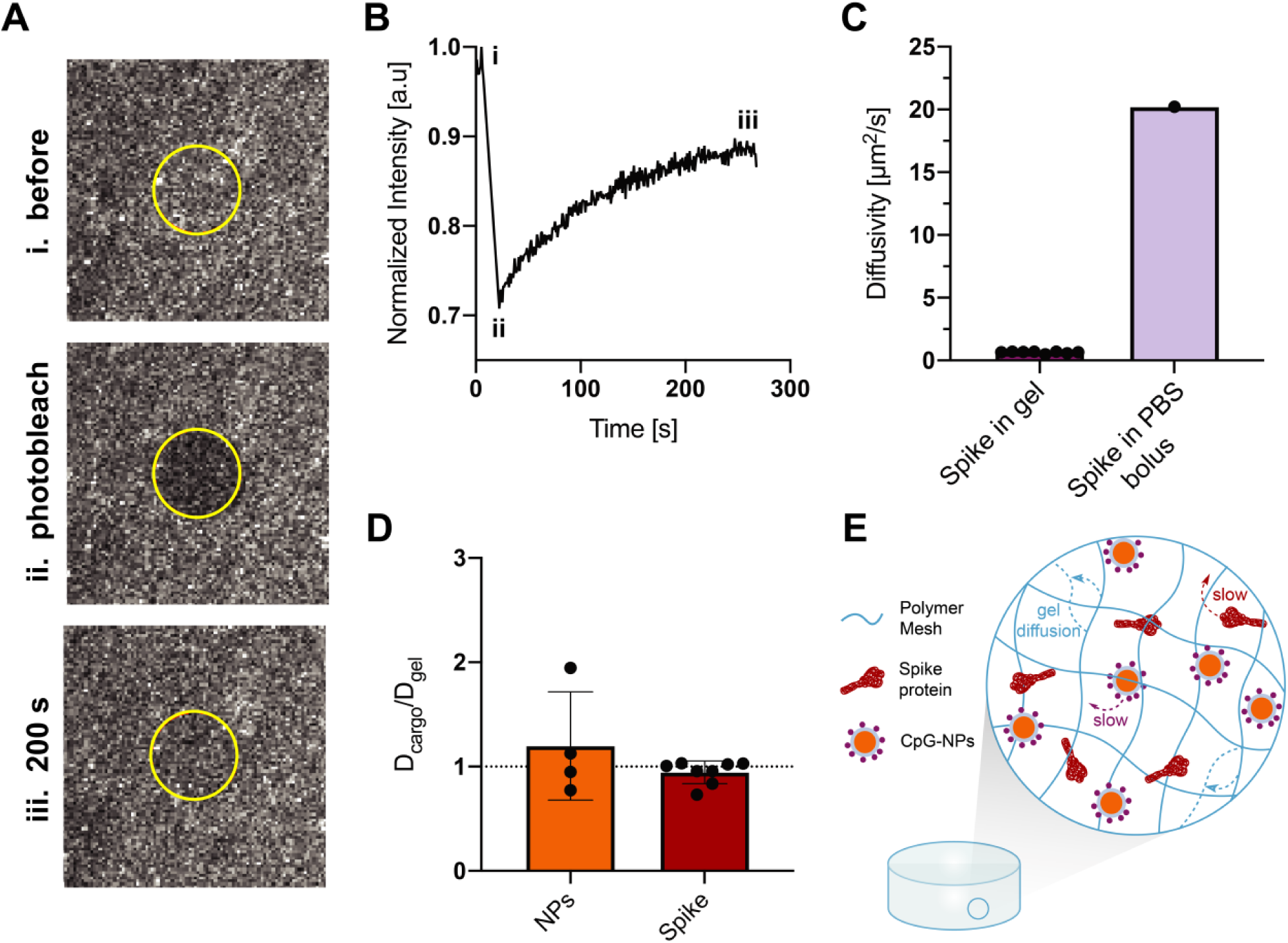
Diffusivity of the cargo and gel components in the CpG-NP hydrogel. **(A)** FRAP microscopy images of the selected area to be photobleached (i) before bleaching, (ii) right after the bleaching process and (iii) after complete fluorescence recovery. **(B)** Representative fluorescence recovery curve over time of the spike protein at a concentration of 0.27 mg/mL of hydrogel. Timepoints representing (A) are outlined on the curve. **(C)** Diffusivities of spike protein in PNP hydrogels (n = 8) measured via FRAP and diffusivity of spike in PBS 1X calculated using Stokes-Einstein equation (Eq. (2)). **(D)** PEG-*b-*PLA NPs and spike protein diffusivities in the hydrogel are measured via FRAP and are represented normalized by D_gel_, the polymer matrix diffusivity. Values close to 1 represent diffusivities similar to the polymer matrix and support the assumption that NPs and spike antigen are caught in the hydrogel network. The dotted line shows D_cargo_/D_gel_ = 1 (n = 4-8). **(E)** Representative schematic of the vaccine loaded PNP hydrogel, showing all the components diffuse slowly within the hydrogel network. All the results are given as mean ± s.d.

We fluorescently labeled both of the main PNP hydrogel components, NPs and HPMC-C_12_, as well as the spike protein antigen. The diffusivity of each component was assessed in distinct experiments to isolate the individual diffusivity effects. From the fluorescence recovery behavior of these molecules, we determined diffusivities, *D,* of hydrogel structural components (HPMC-C_12_), CpG-NPs, and spike protein (n.b. using Eq. 1). FRAP measurements showed that the hydrogel network dramatically reduced the cargo diffusivity of the spike protein by over 30-fold, with a measured diffusivity of *D*_spike_ = 0.64 μm^2^/s, compared to the antigen’s free diffusivity in PBS bolus, which was determined to be *D* = 20.22 μm^2^/s by DLS (n.b. using Eq. 2; Figure 5C). Moreover, the self-diffusion of the PNP matrix, *D*_gel_, was determined by measuring the diffusivity of the HPMC-C_12_ within the fully formulated hydrogel and found to be *D*_gel_ = 0.98 μm^2^/s. The diffusivity of CpG-NPs (*D*_NP_ = 0.68 μm^2^/s) and spike protein in the hydrogel (*D*_spike_ = 0.64 μm^2^/s) were very similar to the self-diffusion of the PNP matrix, resulting in a diffusivity ratio (*D*_cargo_/*D*_gel_) close to 1 for both components (Figure 5D). These results indicate that spike and CpG-NPs, despite their physiochemical differences, are immobilized by the hydrogel’s polymeric network and are diffusing at rates limited by the self-diffusivity of the hydrogel matrix, which arises due to the continuous rearrangement of the dynamic physical PNP network bonds.

### In vivo pharmacokinetic study of spike protein and CpG-NPs in bolus and hydrogel formulations

We further validated the FRAP measurements by evaluating persistence of both spike antigen and CpG-NPs within the hydrogel depot at the injection site. Vaccines containing 10 μg of AF790-spike protein and 20 μg Cy5-CpG equivalent of Cy5-CpG-NPs formulated in either a standard PBS bolus formulation or embedded in hydrogels were subcutaneously injected in SKH1E mice (n=5). Depot formation and persistence at the site of injection was assessed by utilizing brightfield photographic images acquired with a standard camera combined with fluorescent images collected over 16 days from an in vivo Imaging System (IVIS, Figures 6A-6D). AF790-spike in a standard PBS vehicle was nearly undetectable by the end of the first week, while a prolonged signal was observed over two weeks when entrapped in PNP hydrogels (Figure 6A). We measured an enhanced AF790-spike persistence half-life of ∼9 days in PNP hydrogels compared to ∼3.5 days in standard PBS bolus (Figure 6B, 6E). Similarly, the half-life of Cy5-CpG-NP persistence was significantly increased in PNP hydrogels (t_1/2_ >12 days) compared to standard PBS bolus (t_1/2_ ∼0.2 days; Figures 6C-D, 6F). When compared with the previously reported half-life of release for soluble CpG from PNP hydrogels of only ∼2.5 days,^63, 78, 79^ we demonstrated that conjugating CpG onto the PEG-*b*-PLA NPs to form CpG-NPs results in a nearly 5-fold increase in the half-life of release of CpG from the hydrogel depot. Finally, we calculated the ratio of release half-lives for spike protein to CpG-NPs to evaluate the co-release of these two distinct components from the PNP hydrogel depot. While a ratio of over 17 was observed in PBS bolus on account of the large difference in the physicochemical properties of these two species, a ratio of 0.7 was measured when these cargoes are entrapped in PNP hydrogels (Figure 6G). In line with the FRAP data, the significant prolongation in the timeframe of cargo release that aligns with the time-frame of hydrogel erosion previously reported,^76^ coupled with a ratio of release rates for the two cargoes close to 1, further demonstrated that both cargoes are immobilized by the hydrogel’s polymeric network and co-released alongside the erosion of the hydrogel.

**Figure 6:**
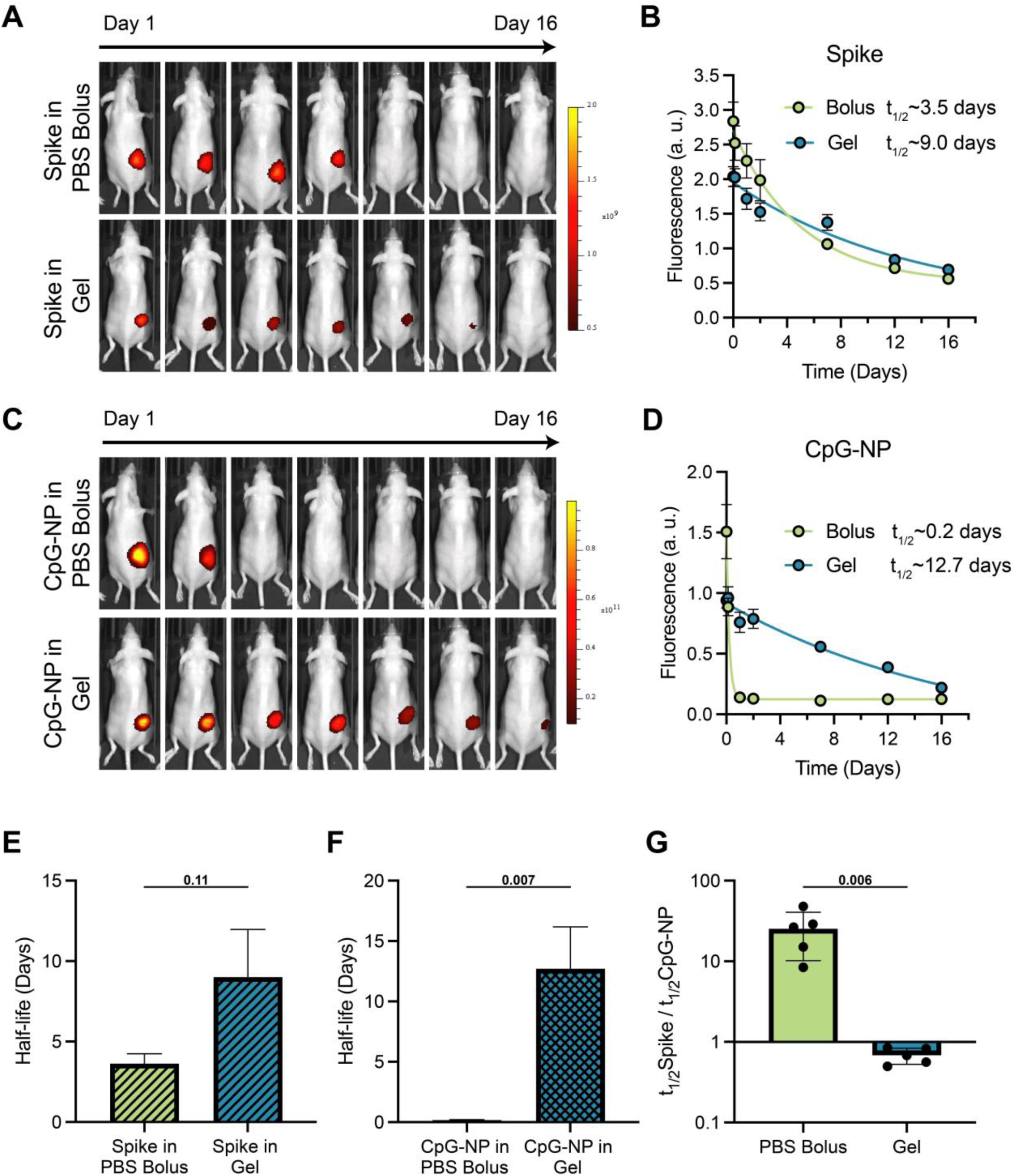
In vivo kinetics of spike and CpG-NP. Mice were immunized with vaccines formulated with Alexa-fluor 790-labeled spike antigen and Cy5-CpG-NP in either PNP hydrogel or PBS 1X bolus formulation. **(A)** Representative images showing the different duration of release of spike protein given as a bolus or hydrogel subcutaneous immunization over 16 days. **(B)** Fluorescent signal from Alexa-fluor 790-labeled spike protein shown in **(A)**. **(C)** Representative images demontrating the different duration of release of CpG-NP given as a bolus or gel subcutaneous immunization over 16 days. **(D)** Fluorescent signal from Alexa-fluor 790-labeled spike protein shown in **(B)**. Images and signal were determined by an In Vivo Imaging System, results are shown as mean ± s.d. (n=5). *p* values listed were determined using unpaired two-tailed t-tests.

### Immunogenicity of COVID-19 vaccines comprising CpG-NP adjuvants

We next investigated the immunogenicity of COVID-19 vaccines comprising spike protein antigen and either soluble CpG or CpG-NPs adjuvants, as well as CpG-NP hydrogels. The spike protein has been one of the most promising antigens used in SARS-CoV-2 subunit vaccine candidates and forms the basis for most clinical vaccines. The spike protein contains the receptor binding domain (RBD) that recognizes the cell surface receptor ACE2, and which is vital for viral fusion and infection. In these experiments, we subcutaneously immunized C57BL/6 mice with vaccines containing 10 μg of spike antigen and adjuvanted with either soluble CpG (20 μg), CpG-NPs (containing an equivalent of 20 μg of CpG), or PEG-*b*-PLA NPs as a vehicle control. These soluble vaccine groups received a prime immunization at week 0 and a boost immunization on week 3, with sera collected weekly from weeks 0-10 (Figure 7A). As we have previously demonstrated robust humoral responses with a single immunization of SARS-CoV-2 RBD hydrogel vaccines,^63, 80^ we also evaluated a single subcutaneous immunization of PNP hydrogels comprising spike antigen (20 μg) adjuvanted with either soluble CpG or CpG-NP (containing an equivalent of 40 μg of CpG) on week 0 (referred as CpG gel and CpG-NP gel, respectively). As with the prime-boost soluble vaccines, sera were collected weekly at weeks 0-10. Thus, these single immunization PNP hydrogel vaccines contained the same total dose of antigen and adjuvant as the complete prime-boost soluble vaccine regimens.

**Figure 7:**
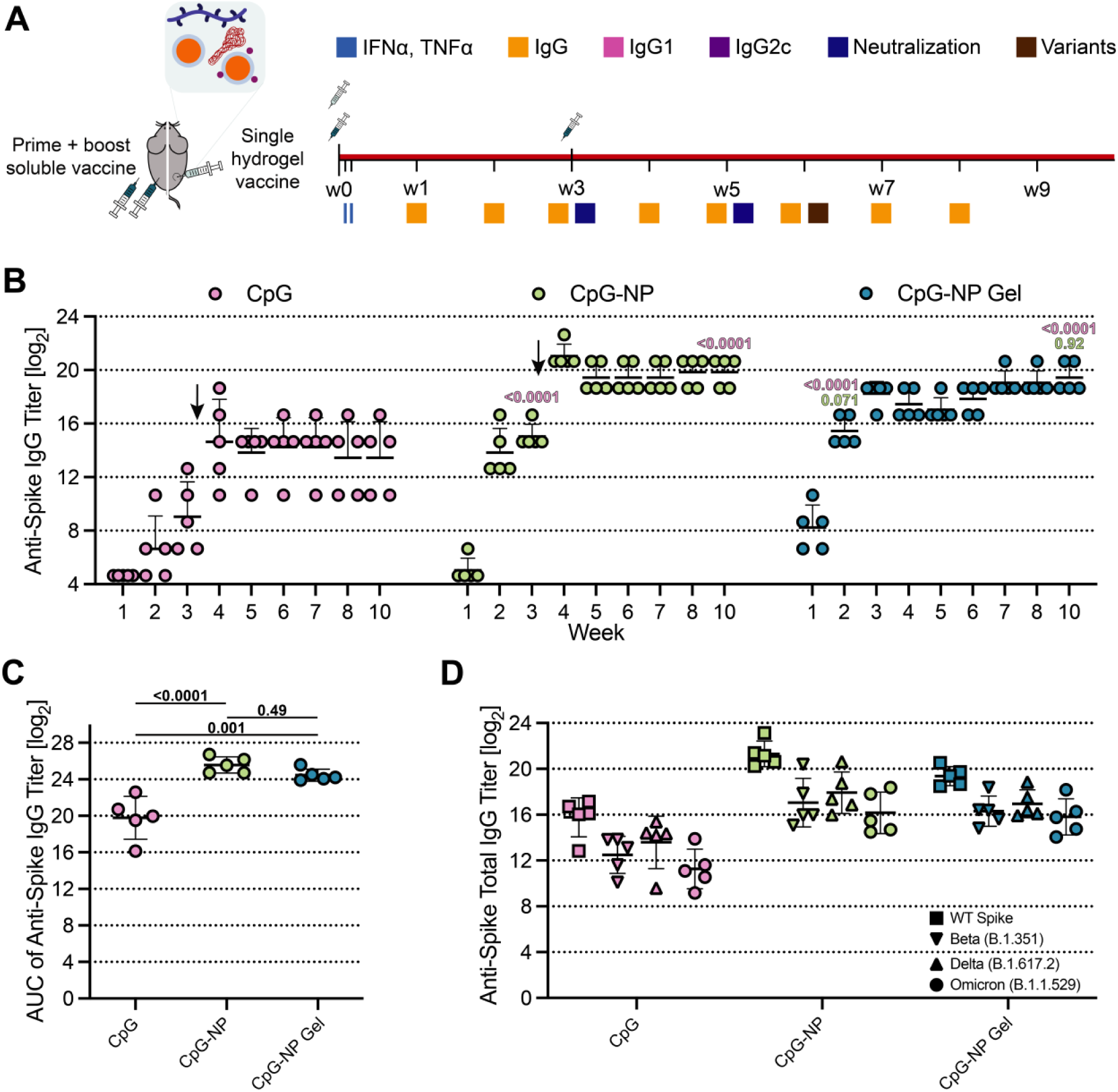
In vivo humoral response to COVID-19 subunit vaccine. **(A)** Timeline of mouse immunizations and blood collection for different assays. Soluble vaccine groups were immunized with a prime dose of 10 µg spike antigen and 20 µg CpG NPs or soluble CpG at day 0 and received a booster injection of the same treatment at day 21. CpG-NP hydrogel group was immunized with a single dose of 20 µg of spike antigen and 40 µg of CpG-NP adjuvant at day 0. Serum was collected over time to determine cytokine levels and IgG titers. IgG1, IgG2b, and IgG2c titers were quantified and neutralization assays were conducted on day 21 and day 35 serum. (**B)** Anti-spike total IgG ELISA endpoint titer of soluble vaccines before and after boosting (arrow) and single immunization CpG-NP hydrogel. **(C)** Area under the curve (AUC) of anti-spike titers from (B). **(D)** Anti-spike IgG ELISA titers from serum collected on week 6, 3 weeks after boosting the soluble vaccine groups.Titers were determined for Wildtype spike as well as Beta (B.1.351), Delta (B.1.617.2, and Omicron (B.1.1.529) variants of the spike protein. Each point represents an individual mouse (n = 5). Data are shown as mean +/- s.d. *p* values listed were determined using a 1way or 2way ANOVA with Tukey’s multiple comparisons test on the logged titer values for IgG titer comparisons (including total IgG and spike variants). *p* values for comparisons are shown above the data points.

High systemic levels of inflammatory cytokines are associated with toxicity in both rodents and humans.^81, 82^ We therefore assessed inflammatory cytokines IFN-α (Figure S7A) and TNF-α (Figure S7B) at 3 h and 24 h after immunization to ensure the vaccines did not drive systemic cytokine responses posing a potential safety risk. No increase in systemic cytokine levels were detected (< 20 pg/mL) across all treatments, including PNP hydrogel immunizations that contained twice the dose of vaccine components. These levels of systematic cytokines remain below previously reported values, suggesting thereby that all treatments were well tolerated.^39, 71, 83^

To assess humoral immune responses, spike-specific immunoglobulin G (IgG) endpoint antibody titers were quantified weekly throughout the experiment. One week post-prime immunization, the endpoint titers were below the detection limit for all of the soluble vaccine groups (except for one mouse in the CpG-NP group) but were sufficiently high in the CpG-NP hydrogel group to suggest all animals had seroconverted (Figures 7B, S8B, *p* < 0.0001 for comparison of hydrogel group to all other groups). This observation is consistent with previous findings where sustained vaccine exposure with PNP hydrogels led to more rapid seroconversion from IgM to IgG, which implies quicker disease protection that is highly desirable in a rapidly-evolving pandemic setting.^62, 63^ CpG-NP hydrogel vaccines elicited higher anti-spike IgG endpoint titers than all soluble vaccines over the first three weeks prior to boost immunization.

Following boost immunization, the CpG-NP group elicited endpoint titers nearly 2 orders of magnitude higher than those elicited by the soluble CpG group (*p* < 0.001 for all time points post-boost). We also observed significantly higher titers for the CpG-NP group compared to all other control groups (with *p* < 0.05 for all time points post-boost). Similarly, the prime-only CpG-NP hydrogel vaccine group, but not unconjugated CpG in PNP hydrogel, elicited comparable endpoint titers to the prime-boost CpG-NP soluble vaccine group from week 3 to the end of the study (*p* > 0.05 at all timepoints; *p* = 0.99 on D70). Notably, significantly increased area under the curve (AUC) of endpoint titers was observed over the entire study period for both the prime-boost CpG-NP soluble group and the CpG-NP hydrogel group compared to unconjugated CpG soluble group, and the controls consisting of unconjugated CpG in PNP hydrogels groups (Figures 7C, S8C). Additionally, consistent with our previous findings, we also observed smaller variability in titer across animals for the prime-only CpG-NP hydrogel group compared to all prime-boost soluble vaccine groups evaluated, which is an important characteristic for ensuring sufficient immunity among all members of a broad population. Overall, these findings demonstrate that CpG-NP adjuvants significantly improve humoral immunity of spike-based vaccines compared to soluble CpG adjuvants, and that sustained co-delivery of CpG-NP adjuvants and spike antigen within PNP hydrogels allows for prime-only single immunization with similarly improved humoral immune responses.

In addition to evaluating humoral responses to the homologous wildtype (WT) SARS-CoV-2 spike variant, we assessed whether the newly designed vaccines can generate broad protection against previously reported SARS-CoV-2 variants of concern such as Beta (B.1.351), Delta (B.1.617.2), and Omicron (B.1.1.529) variants by determining the total IgG endpoint titers against these variants. Across all vaccine groups, decreased titers against all three variants of concern were observed, which is consistent with the known immune-evasion of these variants.^84^ Yet, both prime-boost CpG-NP soluble vaccines and prime-only CpG-NP hydrogel vaccines elicited significantly higher endpoint titers against these three variants of concern compared to prime-boost unconjugated CpG soluble vaccines, prime-boost PEG-*b*-PLA NP hydrogel control vaccines, and prime-only unconjugated CpG hydrogel vaccines (Figures 7D, S8D; *p* < 0.05 for all comparisons). Notably, the endpoint titers against the three variants elicited by prime-boost CpG-NP soluble vaccines and prime-only CpG-NP hydrogel vaccines remained higher than WT endpoint titers produced by the prime-boost unconjugated CpG soluble vaccines. Specifically, CpG-NP soluble and CpG-NP hydrogel vaccines elicited anti-Omicron titers equal to the anti-WT titers elicited by the unconjugated CpG soluble control vaccines (*p* > 0.999 for comparison of both CpG-NP soluble group and CpG-NP hydrogel group’s anti-Omicron titers to unconjugated CpG soluble group’s anti-WT titers). Moreover, we observed a 28% and 24% drop in anti-Omicron titers for prime-boost unconjugated CpG and CpG-NP soluble groups compared to each group’s anti-WT titers, respectively (Figure S9). These decreases were greater than that determined for the CpG-NP hydrogel group, which demonstrated only a 19% drop in anti-Omicron titers compared to anti-WT titers (*p* = 0.16 and *p* = 0.67 for the comparison of the titer reduction of the CpG-NP hydrogel group compared to the CpG-NP and the unconjugated CpG soluble groups, respectively). In sum, both CpG-NP and CpG-NP hydrogel vaccines demonstrated enhanced breadth of humoral immunity against SARS-CoV-2 variants of concern than CpG soluble vaccines.

We next evaluated IgG isotypes at week 7 of the study to assess immune signaling and antibody class-switching following each vaccination. We were especially interested in determining the elicitation of IgG1 and IgG2c antibody responses as these are respectively associated with Th2- and Th1-dominated immune responses.^85^ We found both CpG-NP and CpG-NP hydrogel vaccines exhibited elevated IgG1 endpoint titers compared to other groups (Figures 8A, S10A, *p* < 0.05). Moreover, all groups containing CpG showed elevated IgG2c endpoint titers compared to the PEG-*b*-PLA NP hydrogel control group (Figures 8B, S10B). When assessing the ratio of IgG2c to IgG1, all CpG containing groups were found to elicit an IgG2c/IgG1 ratio close to 1, suggesting balanced Th1 and Th2 responses (Figures 8C, S10C). This observation is consistent with reported studies comparing CpG to other clinical adjuvants such as Alum.^86–88^ Despite previously observing generally more Th2-skewed responses in PNP hydrogels compared to soluble vaccine counterparts,^50, 62, 63^ hydrogels maintained a more balanced response. In the context of COVID-19 infection, clinical studies have found that a rapid onset of a Th1 response resulted in less severe disease outcomes, whereas Th2-skewed responses were associated with greater lung inflammation and higher patient mortality.^89, 90^ The role of the CpG-NP adjuvants in inducing potent Th1 responses may be especially advantageous as a COVID-19 vaccine adjuvant. Future studies will reveal the degree to which CpG-NP adjuvants impact cell-mediated responses, including induction of antigen-specific cytotoxic CD8+ T cells.

**Figure 8:**
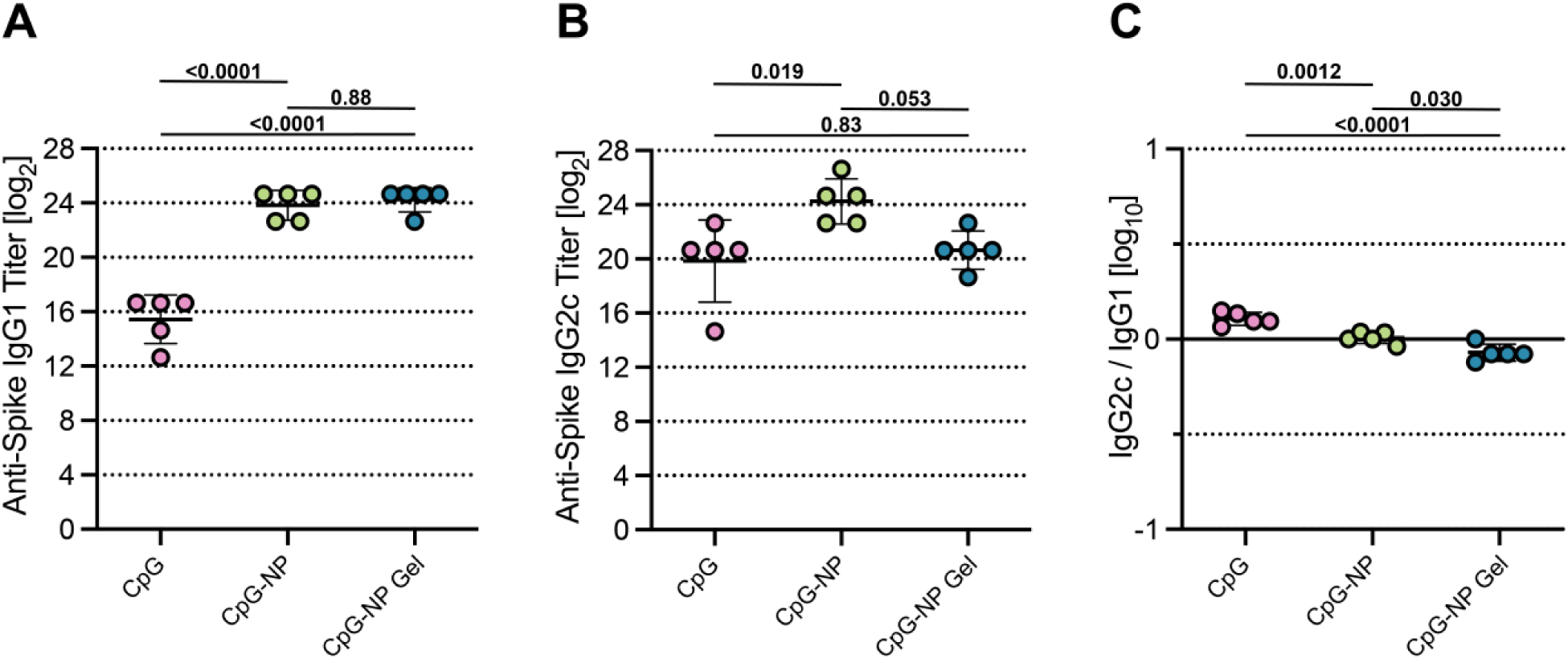
Antibody subtype response to COVID-19 subunit vaccine. Anti-spike IgG1 **(A)** and IgG2c **(B)** titers from serum collected on week 5, 2 weeks after boosting the soluble vaccine groups. **(C)** The ratio of Anti-spike IgG2c to IgG1 post-boost titers. Lower values (below 1) suggest a Th2 response or humoral response, and higher values (above 1) suggest a Th1 response or cellular response. Each point represents an individual mouse (n = 5). Data are shown as mean +/- s.d. *p* values listed were determined using a 1way or ANOVA with Tukey’s multiple comparisons test on the logged titer values for IgG titer comparisons. *p* values for comparisons are shown above the data points.

### SARS-CoV-2 Spike-Pseudotyped Viral Neutralization Assay

We also sought to evaluate the neutralizing activity of the sera from each vaccine group using lentivirus pseudotyped with SARS-CoV-2 spike and determining the inhibition of viral entry into HeLa cells overexpressing the angiotensin-converting enzyme 2 (ACE2) surface receptor (Figure 7A). We first measured the neutralizing activity of sera at week 3 of the study (pre-boost for soluble vaccines) at a single serum dilution of 1:50 (Figures 9A, S11A). Sera from mice immunized with soluble vaccines were found to have at least 50% infectivity, with sera from the PEG-*b*-PLA NP hydrogel control and unconjugated CpG soluble vaccine groups having negligible effect on viral infectivity. On the other hand, sera from mice immunized with CpG-NP hydrogels protected cells from infection (*p* < 0.0001 for comparison of infectivity of CpG-NP hydrogel to all other vaccine groups). This finding suggests that the CpG-NP hydrogel vaccines rapidly generate robust neutralizing activity following a single immunization.

**Figure 9:**
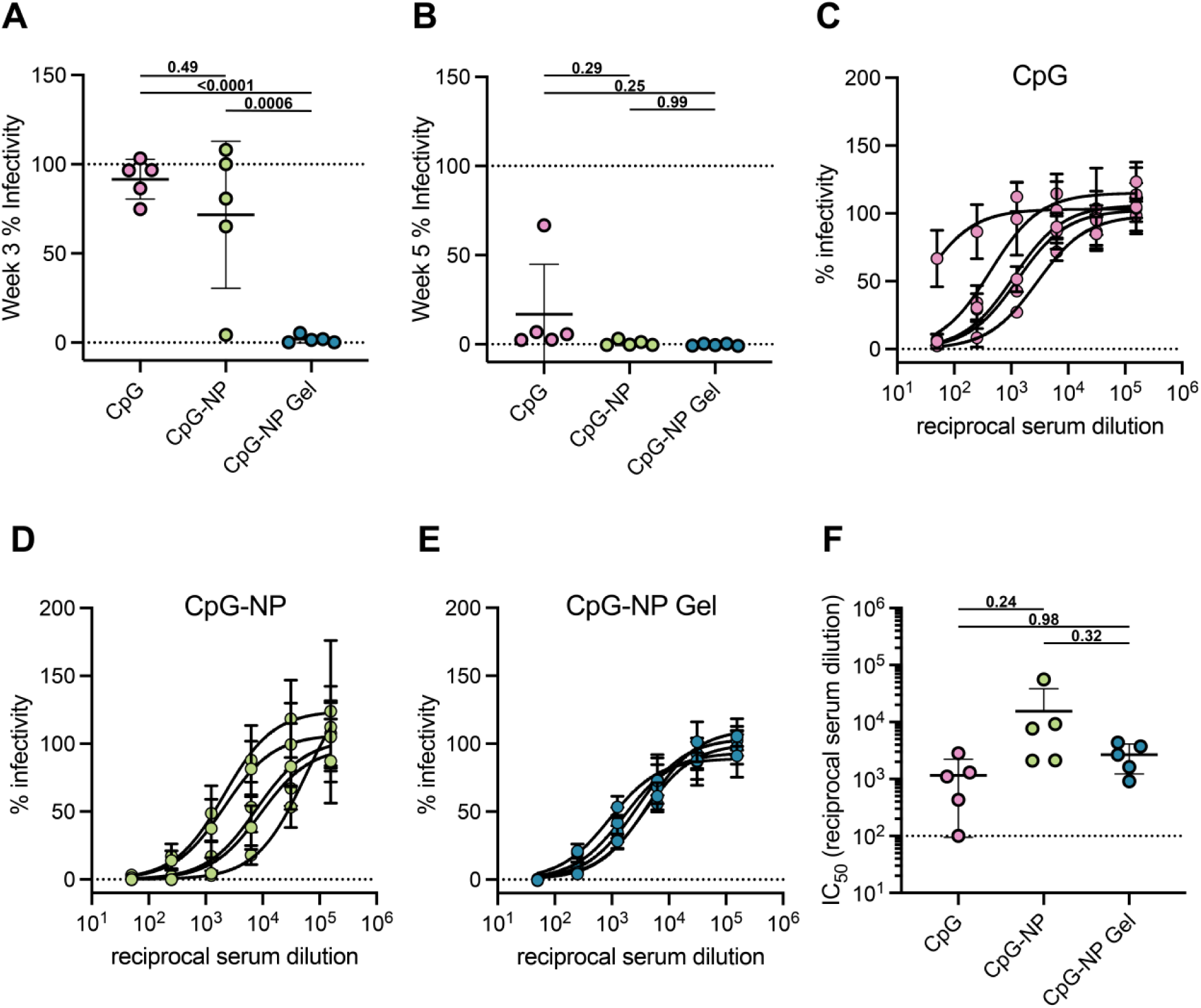
Single immunization of CpG-NP hydrogel elicits neutralizing antibodies in mice. **(A)** Pre-boost (Day 21) spike-pseudotyped viral neutralization assays for the CpG-adjuvanted COVID-19 spike vaccines at a serum dilution of 1:50. **(B)** Post-boost of soluble vaccines (Day 35) spike-pseudotyped viral neutralization assays for the CpG-adjuvanted COVID-19 spike vaccines at a serum dilution of 1:50. **(C-E)** Percent infectivity for all treatment groups at a range of Week 5 serum dilutions as determined by a SARS-CoV-2 spike-pseudotyped viral neutralization assay. **(F)** Comparison of IC_50_ values determined from neutralization curves on Day 35 for soluble vaccine formulations (prime-boosted) and hydrogel vaccine (single immunization) following immunization with CpG-adjuvanted COVID-19 spike vaccines. Each data point represents an individual mouse (n = 5). Data are shown as mean +/- s.d. *p* values listed were determined using a 1way ANOVA with Tukey’s multiple comparisons. *p* values for comparisons are shown above the data points.

We then measured the neutralizing activity of sera at week 5 of the study (two weeks post-boost for soluble vaccines) from all vaccine formulations at a single dilution of 1:50 (Figures 9B, S11B). We determined that sera from soluble vaccine groups resulted in an infectivity less than 50% only after boosting, while sera from the single-immunization CpG-NP hydrogel vaccine group exhibited protection against infection. We then assayed a range of sera concentrations from all groups to determine the half maximal inhibition of infectivity (IC_50_) (Figures 9C-F, S11C-D). The prime-boost CpG-NP soluble vaccine group was found to have the most potent neutralization (IC_50_ ∼1.5×10^4^), followed by the prime-only CpG-NP hydrogel group (IC_50_ ∼ 2.7×10^3^). Even though the soluble vaccine groups had just received a booster vaccination 2 weeks prior, the measured IC_50_ from the single-immunization CpG-NP hydrogel group was comparable to the other prime-boost soluble vaccine groups (*p* > 0.05 comparing all groups to the CpG-NP hydrogel group).

Overall, we determined that a single immunization of CpG-NP hydrogel vaccines was similarly effective as a prime-boost immunization regimen with soluble CpG-NP vaccines in terms of overall antibody titer, durability of titer responses, breadth of antibody responses, balanced Th1 and Th2 responses, as well as neutralization activity. Further, vaccines formulated with CpG-NPs (e.g., both prime-only CpG-NP hydrogel and prime-boost CpG-NP soluble vaccines) demonstrated superior overall humoral responses compared to vaccines comprising soluble CpG as an adjuvant.

## CONCLUSIONS

In conclusion, we have developed a potent adjuvant nanoparticle platform enabling the presentation of TLR9 agonists on the surface of PEG-*b*-PLA NPs. Our facile synthetic and formulation approach allows for precise control of the valency of adjuvant distribution on the NPs surface. We showed that the density of CpG presentation strongly influenced the activation of TLR9, and that an intermediate density of CpG on the NP surface (e.g., 30% CpG-NPs) exhibited the greatest potency *in vitro* compared to soluble CpG. When these CpG-NPs were used as adjuvants in candidate COVID-19 vaccines using the SARS-CoV-2 spike protein, we found that they elicited superior humoral responses compared to soluble CpG adjuvants. Indeed, vaccines comprising CpG-NP adjuvants elicited more potent and sustained antibody titers, more robust breath of recognition of immune-evading variants of concern, balanced Th1 to Th2 responses, and more strongly neutralizing antibody responses than soluble CpG. These promising CpG-NP adjuvants were further evaluated within PNP hydrogels to enhance the spatiotemporal control of vaccine delivery. Embedding of CpG-NPs within PNP hydrogels was found to negligibly impact the rheological properties compared to standard PNP hydrogels. Additionally, we confirmed the immobilization of both CpG-NPs and SARS-CoV-2 spike protein in the hydrogel’s network results in similar diffusive properties, despite their physicochemical differences, thereby enabling sustained co-delivery of both vaccine components. Importantly, a single immunization of CpG-NP hydrogels generated comparable humoral responses to a prime-boost regimen of CpG-NP soluble vaccines. The promising results of single-immunization CpG-NP hydrogel vaccines could reduce clinical vaccination costs, increase patient compliance, and ultimately result in more rapid uptake of vaccines and higher vaccination rates, which are all key elements when fighting against a rapidly evolving pandemic.

## EXPERIMENTAL SECTION

### Materials

Poly(ethylene glycol)methyl ether 5000 Da (PEG-methyl ether), Poly(ethylene glycol)α-hydroxy-ω-azido terminated 5000 Da (N_3_-PEG-OH), 3,6-Dimethyl-1,4-dioxane-2,5-dione (Lactide), 1,8-diazabicyclo(5.4.0)undec-7-ene (DBU, 98%), (Hydroxypropyl)methyl cellulose (HPMC, meets USP testing specifications), *N,N*-Diisopropylethylamine (Hunig’s base), *N*-methyl-2-pyrrolidone (NMP), 1-dodecyl isocyanate (99%), mini Quick Spin Oligo columns (Sephadex G-25 Superfine packing material), Sepharose CL-6B crosslinked, and bovine serum albumin (BSA) were purchased from Sigma-Aldrich. CpG-C 2395 (5’-TCGTCGTTTTCGGCGCGCGCCG-3’) oligonucleotide, and Aluminum hydroxide gel (Alhydrogel adjuvant 2%), zeocin were purchased from InvivoGen. Amino CpG-C 2395 (5’Amino Modifier C_6_, 5’-NH_2_-TCGTCGTTTTCGGCGCGCGCCG-3’) and Amino CpG-C 2395 Cyanine 5 (5’Amino Modifier C_6_, 5’-NH_2_-TCGTCGTTTTCGGCGCGCGCCG-3’Cy5Sp) were purchased from Integrated DNA Technologies (IDT). Dibenzocyclooctyne-PEG4-*N*-hydroxysuccinimidyl ester (DBCO-PEG4-NHS ester) and Alexa Fluor 647 DBCO (AFDye 647 DBCO) were purchased from Click Chemistry Tools. Alexa Fluor 790 Succinimidyl Ester, Dulbecco’s modified Eagle’s medium (DMEM, Gibco), phosphate buffered saline (PBS pH 7.4, Gibco), Invitrogen E-Gel EX Agarose Gels 4%, and DAPI (4’,6-Diamidino-2-Phenylindole, Dihydrochloride, D1306) were purchased from Thermo Fisher Scientific. Heat Inactivated fetal bovine serum (HI-FBS) was purchased from Atlanta Biologicals. IFN-α cytokine enzyme-linked immunosorbent assay (ELISA) kit was purchased from PBL Assay Science, and TNF-α cytokine ELISA kit was purchased from R&D Systems (Fisher Scientific). Goat anti-mouse IgG Fc secondary antibody (A16084) HRP (Horseradish peroxidase) was purchased from Invitrogen. Alexa Fluor 488 Anti-alpha 1 Sodium Potassium ATPase antibody (ab197496), Goat anti-mouse IgG1 and IgG2c Fc secondary antibodies (ab97250, ab97255) HRP were purchased from Abcam. 3,3’,5,5’-Tetramethylbenzidine (TMB) ELISA Substrate, high sensitivity was acquired from Abcam. HIS Lite Cy3 Bis NTA-Ni Complex was purchased from AAT Bioquest. Unless otherwise stated, all chemicals were used as received without further purification.

### Synthesis of PEG-*b-*PLA

PEG-*b*-PLA was prepared as previously reported.^58^ Prior to use, commercial lactide was recrystallized in ethyl acetate and Dichloromethane (DCM) was dried via cryo distillation. Under inert atmosphere (N_2_), PEG-methyl ether (5 kDa, 0.25 g, 4.1 mmol) and DBU (15 µL, 0.1 mmol) were dissolved in 1 mL of anhydrous DCM. Lactide (1.0 g, 6.9 mmol) was dissolved under N_2_ in 3 mL of anhydrous DCM. The lactide solution was then quickly added to the PEG/DBU mixture and was allowed to polymerize for 8 min at room temperature. The reaction was then quenched with an acetic acid aqueous solution and the polymer precipitated into a 1:1 mixture of ethyl ether and hexanes, collected by centrifugation and dried under vacuum. NMR spectroscopic data, Mn and Dispersity were in agreement with those previously described.

### Synthesis of azide PEG-*b-*PLA

Azide-PEG-b-PLA was synthesized according to the literature.^65, 75^ Prior to use, DCM was dried using 3-4 Å molecular sieves and N_3_-PEG-OH was dried under vacuum overnight. Under inert atmosphere, a solution of N_3_-PEG-OH (0.5 g, 5 kDa, 100 µmol) and DBU (30 µL, 9.29 mmol) in anhydrous DCM (1 mL) was rapidly added to a solution of lactide (2.0 g, 13.9 mmol) in anhydrous DCM (10 mL) and stirred for 8 min at room temperature. The reaction mixture was quenched with an acetic acid aqueous solution, precipitated in a mixture of ethyl ether: hexanes (1:1), centrifuged and dried under vacuum overnight. NMR spectroscopic data, Mn and Dispersity were in agreement with those previously described.

### Synthesis of DBCO-CpG intermediate

DBCO-PEG4-NHS ester (3.78 mg, 5.8 µmol) was dissolved in DMSO (40 µL). The solution was diluted with PBS 1X to reach a final concentration of 10 mM. NH_2_-CpG or NH_2_-CpG-Cy5 (0.58 µmol) was then reacted with DBCO-PEG4-NHS ester solution for 6 h at room temperature. The solution was purified by size-exclusion chromatography in PBS 1X using a Sephadex G-25 Superfine (mini Quick Spin Oligo) column and stored at −20 °C.

### NPs formulation and conjugation

PEG-*b*-PLA NPs were prepared as previously described.^50, 64^ A 1 mL solution of PEG-*b*-PLA and N_3_-PEG-*b*-PLA in 75:25 ACN:DMSO (50 mg/mL) was added dropwise to 10 mL of Milli-Q water stirring at 600 rpm. The particles solution was purified in centrifugal filters (Amicon Ultra, MWCO 10 kDa) at 4500 RCF for 1 h and resuspended in PBS 1X to reach a final concentration of 200 mg/mL. DBCO-CpG or DBCO-CpG-Cy5 (3 eq) and N_3_-PEG-*b*-PLA NPs (1 eq) were reacted *via* copper-free click chemistry in PBS 1X for 12 h at room temperature. After reaction completion, CpG conjugated NPs were purified by size-exclusion chromatography on a Sepharose CL-6B matrix eluting with PBS 1X. Successful purification of the CpG-NPs from unreacted soluble CpG was confirmed *via* aqueous SEC measurements and agarose gel electrophoresis (4%). The CpG concentration on the NPs was determined through absorption calibration curves at 280 nm acquired using a Synergy H1 Microplate Reader (BioTek Instruments). An individual calibration curve for each NPs valency and TLR9 agonist class was recorded. Conversions between 88 and 97% were measured.

### HPMC-C_12_ synthesis

HPMC-C_12_ was prepared according to a previously reported procedure.^58^ HPMC (1.0 g) was dissolved in anhydrous NMP (45 mL) by stirring at 80 °C for 1 h. Once cooled to room temperature, 1-dodecyl isocyanate (105 mg, 0.5 mmol) and Hunig’s base, acting as the catalyst (∼ 3 drops) were dissolved in 5 mL of anhydrous NMP. This solution was then added dropwise to the reaction mixture, which was stirred at room temperature for 16 h. The polymer was precipitated using acetone, redissolved in Milli-Q water (∼ 2 wt%) and dialyzed (3 kDa MWCO) against water for 4 days. The polymer mixture was then lyophilized and reconstituted to a 60 mg/mL solution in sterile PBS 1X.

### DMF-SEC measurements

Apparent molecular weight and dispersity were obtained after passing through two size exclusion chromatography columns (Resolve Mixed Bed Low DVB, inner diameter ID of 7.8 mm, M_w_ range: 200–600,000 g/mol, Jordi Labs) in a mobile phase of *N,N*-dimethylformamide (DMF) with 10 mM LiBr at 35 °C and a flow rate of 1.0 mL/min (Dionex Ultimate 3000 pump, degasser, and autosampler, Thermo Fisher Scientific). Before injection, samples at a concentration of 5 mg/mL were filtered through a 0.22 μm nylon membrane.

### Aqueous-SEC measurements

SEC traces were determined after passing through a size-exclusion chromatography column [5000 to 5,000,000 g/mol]; Superose 6 Increase 10/300 GL (GE Healthcare) in a mobile phase of PBS containing 300 parts per million of sodium azide and at a flow rate of 0.75 mL/min (Dionex Ultimate 3000 pump, degasser, and autosampler, Thermo Fisher Scientific). Detection consisted of an Optilab T-rEX (Wyatt Technology Corporation) refractive index detector operating at 658 nm and a diode array detector operating at 280 nm (Dionex Ultimate 3000, Thermo Fischer Scientific). Before injection, samples at a concentration of 1 mg/mL were filtered through a 0.22 µm PVDF membrane.

### Dynamic light scattering and zeta potential measurements

The hydrodynamic diameter and surface charge of the NPs were respectively measured on a DynaPro II plate reader (Wyatt Technology) and a Zetasizer Nano Zs (Malvern Instruments). Three independent measurements were performed for each sample.

### Alexa Fluor 790-conjugated spike protein

A pre-mixed solution of AF-790 Succinimidyl Ester (30 μg, 0.017 µmol, 8 eq, 5 mg/ml stock solution in DMSO) in PBS 1X was added to a solution of Spike protein (300 μg, 0.0021 µmol, 1 eq) in PBS 1X. A volume ratio of dye (1/10) and protein (9/10) was respected. The reaction was conducted in the dark for 4 h at RT with mild shaking. The solution was quenched by diluting 2-fold with PBS 1X and purified in centrifugal filters (Amicon Ultra, 10 kDa MWCO 0.5 mL) at 14 g for 10 min. The purification step was repeated until all excess dye was removed. The solution was then resuspended in PBS 1X and stored at −20C.

### PNP and CpG-NP Hydrogel formulation

CpG polymer-nanoparticle (CpG-NP) hydrogels were formed at 2 wt% HPMC-C_12_ and 10 wt% mixture of PEG-*b*-PLA and CpG-PEG-*b*-PLA NPs in PBS 1X. Hydrogels were prepared by mixing a 3:2:1 weight ratio of 6 wt% HPMC-C12 polymer solution, 20 wt% NPs solution, and PBS 1X. Based on the desired adjuvant dosing, 30% CpG conjugated NPs were mixed with non-conjugated PEG-*b*-PLA NPs prior to hydrogel formation. Hydrogels were formed by mixing the solutions using syringes connected through an elbow mixer.

### Rheological Characterization of PNP hydrogels

Rheological characterization was completed on a Discovery HR-2 Rheometer (TA Instruments). Measurements were performed using a 20 mm serrated plate geometry at 25 °C and at 500 µm gap height. Dynamic oscillatory frequency sweeps were conducted at a constant 1% strain and angular frequencies from 0.1 to 100 rad/s. Amplitude sweeps were performed at a constant angular frequency of 10 rad/s from 0.5% to 10,000% strain. Flow sweep, steady shear experiments were performed at shear rates from 50 to 0.005 1/s, whereas stress-controlled flow sweep measurements were conducted at shear rates from 0.001 to 10 1/s. Step-shear experiments were performed by alternating between low shear rates (0.1 rad/s for 60 s) and high shear rates (10 rad/s for 30 s) for three full cycles. Yield stress values were extrapolated from stress-controlled flow sweep and amplitude sweep measurements.

### FRAP Analysis

Fluorescence recovery after photobleaching (FRAP) was performed on PNP hydrogel and CpG-NP hydrogel formulations using a confocal LSM780 microscope. Each individual component of the hydrogel was labelled with a fluorescent dye and analyzed in separate samples. NP-tethered AF647 (10 wt%), rhodamine-conjugated HPMC-C_12_ (2 wt%), and His-tagged SARS-CoV-2 spike conjugated with HIS-Lite-Cy3 Bis NTA-Ni Complex (0.27 mg per mL of hydrogel) were used to visualize diffusion of the vaccine cargo and hydrogel components. Samples were imaged using low intensity lasers to collect an initial level of fluorescence. Then a high intensity laser with a diameter of 25 µm was focused on the region of interest (ROI) for 10 s to bleach a circular area.

Subsequently, fluorescence emission data was recorded for 4 min to create an exponential fluorescence recovery curve. For each sample, replicate measurements (n = 2-5) were taken at multiple locations. The diffusion coefficient *D* was calculated according to the following equation^91^:

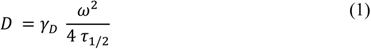

where the constant *γ_D_* = *τ_1/2_*/*τ_D_*, with *τ_1/2_* being the time to half recovery, *τ_D_* the characteristic diffusion time, both yielded by the ZEN software, and *ω* the radius of the bleached ROI. The diffusivity of the SARS-CoV-2 spike protein antigen in PBS 1X was calculated using the Stokes-Einstein law equation for diffusion^79^:

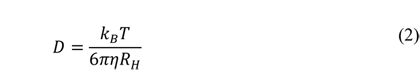

with *k_B_* being the Boltzmann’s constant, *T* the temperature in Kelvin, *η* the solvent viscosity, and *R_H_* the solute hydrodynamic radius. The hydrodynamic radius of the spike protein was measured via DLS to be *R_H_* = 12.2 nm, whereas *η* for PBS 1X was approximated to be 0.8872 mPa s at 25 °C. The measured *R_H_* agrees with the value published in literature and measured via Cryo-EM.^92^

### In vitro reporter assays

The Raw-Blue (NF-kB-SEAP) reporter cell line (Invivogen, raw-sp) and THP-1 hTLR9 reporter cell line (Invivogen, thpd-htlr9) were used to evaluate the effect of the TLR9 agonist valency conjugated to PEG-*b*-PLA NPs. The cells were cultured at 37 °C with 5% CO_2_ in DMEM supplemented with L-glutamine (2 mM), D-glucose (4.5 g/L), 10% HI-FBS, penicillin (100 U/mL), and streptomycin (100 µg) for Raw-Blue cells and in RPMI 1640 supplemented with L-glutamine (2 mM), D-glucose (4.5 g/L), 10% HI-FBS, penicillin (100 U/mL) and streptomycin (100 µg) for THP-1 cells. Every other passage, zeocin (100 µg/mL) and other selective antibiotics were added to the culture medium. Serial dilutions of soluble CpG and different CpG-NP formulations were added to a 96-well tissue culture treated plate to achieve final concentrations between 30 and 3.1 µg/mL of TLR9 agonist. Non-conjugated PEG-*b*-PLA NP was used as a negative control. 100,000 cells were added to each well in 180 µL of media and were incubated for 21 h at 37 °C in a CO_2_ (5%) incubator. Manufacturer instructions were followed for SEAP quantification, and absorbance levels detected at 655 nm after 3 h incubation with QUANTI-Blue Solution (Invivogen). The absorbances of RAW-Blue assay were normalized to absorbance intensity at the highest and lowest dilutions. Normalized nonlinear regression fits were found using the “Log(agonist) vs. response – EC_50_” function in GraphPad Prism 8.4 software. Data normalization and analysis was performed using GraphPad Prism.

### In vitro cellular uptake assays

50,000 Raw-Blue cells were plated on glass dishes (ibidi, 81158) and incubated for 48 hours at 37 °C with 5% CO_2_ in DMEM supplemented with L-glutamine (2 mM), D-glucose (4.5 g/L), 10% HI-FBS, penicillin (100 U/mL), and streptomycin (100 µg). The media was replaced with the same DMEM based media containing soluble CpG, 30% CpG-NP, or 50% CpG-NP at a concentration of 5 μg equivalent of CpG. Cells were co-cultured overnight at 37 °C with 5% CO_2_. The media was then aspirated, and the cells were fixed with 4% PFA in PBS 1X at 37°C with 5% CO_2_ for 15 min before washing with PBS 1X. To stain the cell wall, cells in each glass slide were stained with 400 μL of AF488-ATP antibody (1:100 dilution in 1% BSA in PBS 1X) for 40 min in RT in the dark. Cells were washed 3 times with PBS 1X before incubating with DAPI (300 nM in PBS 1X) for 2 min at RT in the dark. Cells were washed 3 times with PBS 1X before imagining with a confocal microscopy (LSM780).

### Animal studies

Six-to-seven weeks old female C57BL/6 (B6) and SKH1E mice were obtained from Charles River, housed in the animal facility at Stanford University and cared for according to Institutional Animal Care and Use guidelines. All animal studies were performed in accordance with the National Institutes of Health guidelines and the approval of Stanford Administrative Panel on Laboratory Animal Care. The day before vaccine administration, mice were shaved in order to receive a subcutaneous injection of vaccine on the right side of their backs. Mouse blood was collected from the tail vein each week for 10 weeks.

### In vivo biodistribution study of soluble CpG and CpG-NPs

C57BL/6 mice were injected subcutaneously in the right flank with 100 μL of PBS 1X buffer containing 10 μg CpG equivalent of either Cy5-CpG or Cy5-CpG-NPs. Mice were euthanized 3 h post-injection with CO_2_ and their major organs (liver, spleen, kidneys, and ipsilateral lymph nodes) were imaged using an In Vivo Imagining System (IVIS Lago). Imaging procedures and data analysis methods were identical to those thoroughly described in previously published work.^76, 93,^_94_ Cy5-CpG were imaged using auto exposure time, excitation wavelength of 640 nm, and emission wavelength of 670 nm (binning: medium, F/stop: 4).

### In vivo pharmacokinetic study of spike protein and CpG-NPs in bolus and hydrogel formulations

SKH1E mice were immunized subcutaneously in the right flank with 100 μL of soluble or hydrogel vaccines containing 10 μg of AF790-spike protein and 20 μg CpG equivalent of Cy5-CpG NPs. Hydrogels were formulated as described in the previous sections. Mice were imaged over 16 days using the in vivo Imaging System (IVIS Lago). Imaging procedures and data analysis were identical to those performed on PNP hydrogels and described in prior work. ^76, 93, 94^

AF790-Spike proteins were imaged using auto exposure time, excitation wavelength of 780 nm, and emission wavelength of 845 nm (binning: medium, F/stop: 2). Cy5-CpG were imaged using auto exposure time, excitation wavelength of 640 nm, and emission wavelength of 670 nm (binning: medium, F/stop: 4). Average radiant efficiency was quantified. Half-lives of spike protein and CpG retention were obtained by fitting fluorescence intensity values between day 0 and 16 to single phase exponential decay models. Data analysis was performed using GraphPad Prism.

### Vaccine formulation

SARS-CoV-2 spike protein vaccines were injected subcutaneously either in form of a soluble injection or of a hydrogel. For soluble injections, vaccines were formulated in 100 μL of PBS 1X and contained a 10 µg antigen dose of spike S1+S2 ECD (R683A, R685A, F817P, A892P, A899P, A942P, K986P, V987P)-His Recombinant Protein (Sino Biological 40589-V08H4) and a 20 µg 30% CpG-NPs adjuvant dose; boosting was performed on day 21. For the CpG-NP hydrogels, the dose was doubled and contained 20 µg of antigen and 40 µg of CpG-NPs adjuvant formulated in 150 μL of hydrogel; no boosting was performed. Control groups were composed of 100 μL of soluble formulations containing 10 μg of spike protein and non-conjugated PEG-*b*-PLA NPs or soluble CpG (20 µg, IDT) vaccines as well as 150 μL of soluble CpG (40 μg) in PNP hydrogels. Mouse blood was collected from the tail vein each week for 10 weeks. To analyze early cytokine response, blood was collected at 0 h, at 3 h and 24 h from injection and stored at −80 °C. The serum samples were analyzed for IFN-α and TNF-α levels and the concentrations were determined via enzyme-linked immunosorbent assay (ELISA) according to manufacturer’s instructions and were calculated from standard curves. Absorbance was measured with a Synergy H1 microplate reader (BioTek Instruments) at 450 nm.

### Mouse Serum ELISAs

Serum Anti-spike IgG antibody endpoint titers were measured using an ELISA. Maxisorp plates (Thermofisher) were coated with SARS-CoV-2 spike protein (Sino Biological 40591-V08H4), the mutant spike from Beta B.1.351 (Sino Biological 40591-V08H12), the mutant spike from Delta B.1.617.2 (Sino Biological 40591-V08H23), or the mutant spike from Omicron B.1.1.529 (Sino Biological 40591-V08H41) at 2 µg/mL in PBS 1X overnight at 4 °C and subsequently blocked with PBS 1X containing 1 wt% BSA for 1 h at 25 °C. Serum samples were serially diluted and incubated in the coated plates for 2 h at 25 °C, and goat-anti-mouse IgG Fc-HRP (1:10,000), IgG1 Fc-HRP (1:10,000), or IgG2c (1:10,000) was added for 1h at 25 °C. Plates were developed with TMB substrate, the reaction stopped with 1 M HCl and the plates analyzed using a Synergy H1 microplate reader (BioTek Instruments) at 450 nm. End point titers were defined as the highest serum dilution at the one for which an optical density above 0.1 was detected.

### SARS-CoV-2 Spike-pseudotyped Viral Neutralization Assay

Neutralization assays were conducted as previously described.^95^ Briefly, SARS-CoV-2 spike pseudotyped lentivirus was produced in HEK239T cells and cells seeded at six million cells the day prior to transfection. A five-plasmid system was used for viral production. Plasmids were added to filter-sterilized water and HEPES-buffered saline was added dropwise to reach a final volume of 1 mL. To form transfection complexes, CaCl_2_ was added dropwise to the gently agitated solution. The transfection reactions were incubated for 20 min at RT and then added to plated cells. Virus-containing culture supernatants were harvested ∼72 hours after transfection via centrifugation and filtered through a 0.45 µm syringe filter. Viral stocks were stored at −80 °C. For the neutralization assays, ACE2/HeLa cells were plated 1 to 2 days prior to infection and mouse serum was heat inactivated at 56 °C for 30 min prior to use. Mouse serum (1:50 dilution) and virus were diluted in cell culture medium and supplemented with polybrene at a final concentration of 5 µg/mL. Serum/virus solution at 1:50 were incubated at 37 °C for 1 h. After the incubation period, the media was removed from the cells and incubated with the serum/virus solution at 37°C for 48h. After complete incubation, the cells were then lysed using BriteLite (Perkin Elmer) luciferase readout reagent, and luminescence was measured with a BioTek plate reader. Each plate was normalized by averaging the readout from the wells containing only the virus or only the cells.

### Statistical Analysis

All results are expressed as mean ± standard deviation (*s.d*). Comparison between two groups were conducted by a two-tailed Student’s t-test. One-way ANOVA tests with a Tukey’s multiple-comparisons test was used for comparison across multiple groups. For plots displaying multiple time points or protection against different variants, *p* values were determined with a 2way ANOVA with Tukey’s multiple-comparisons test. Statistical analysis was performed using GraphPad Prism 8.4 (GraphPad Software). Statistical significance was considered as *p* < 0.05.

## ASSOCIATED CONTENT

### Supporting Information

The following files are available free of charge.

Additional tables and figures of synthesis, purification, and characterization of CpG-NPs, and additional immune response data and figures. (PDF)

## AUTHOR INFORMATION

### Corresponding Author

Eric A. Appel

Department of Materials Science & Engineering, Department of Bioengineering, Stanford ChEM-H Institute, Department of Pediatrics – Endocrinology, and Woods Institute for the Environment, Stanford University, Stanford CA 94305, USA

E

### Author Contributions

BSO, VTM, ECG and EAA designed the broad concepts and research studies. JB, AEP, OMS, and HLP designed specific experiments. BSO, VTM, JB, ECG, AEP, OMS, JY, AN, and HLP performed research and experiments. BSO, JB, VTM, and EAA wrote the paper. BSO, JB, OSM, and JY edited the paper. The manuscript was written through contributions of all authors. All authors have given approval to the final version of the manuscript. ‡These authors contributed equally.

## ACKNOWLEDGMENTS

We would like to thank all members of the Appel lab for their useful discussion and advice throughout this project as well as the staff of the BioE/ChemE Animal Facility who cared for our mice. This work was financially supported by the Center for Human Systems Immunology with the Bill & Melinda Gates Foundation (OPP1113682; OPP1211043; INV027411), the American Cancer Society (RSG-18-133-01-CDD), the Goldman Sachs Foundation (administered by the Stanford Cancer Institute, SPO# 162509), and a Bio-X Interdisciplinary Initiatives Seed Grant. BSO is grateful for an Eastman Kodak Fellowship. OMS and JY are thankful for a National Science Foundation Graduate Research Fellowship. AN is thankful for the Paul and Mildred Berg Fellowship. This work was also supported by the Stanford Maternal and Child Health Research Institute postdoctoral fellowship (to AEP). We thank Dr. Jesse Bloom, Kate Crawford, Dr. Dennis Burton, and Dr. Deli Huang for sharing the plasmids, cells, and invaluable advice for implementation of the spike-pseudotyped lentiviral neutralization assay (to AEP). We also thank Dr. Peter S. Kim for his advice and support.

