## Supplemental Information for "Nanoparticle-Conjugated TLR9 Agonists Improve the Potency, Durability, and Breadth of COVID-19 Vaccines"

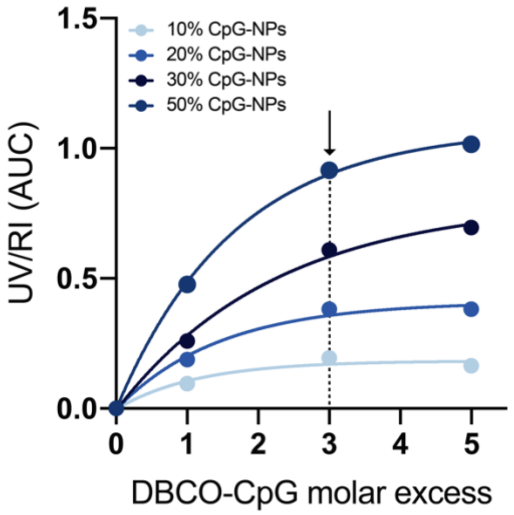


**Figure S1:** Influence of DBCO-CpG molar excess on the click reaction conversion. Three equivalents of DBCO-CpG result in reaction conversions higher than 90%.

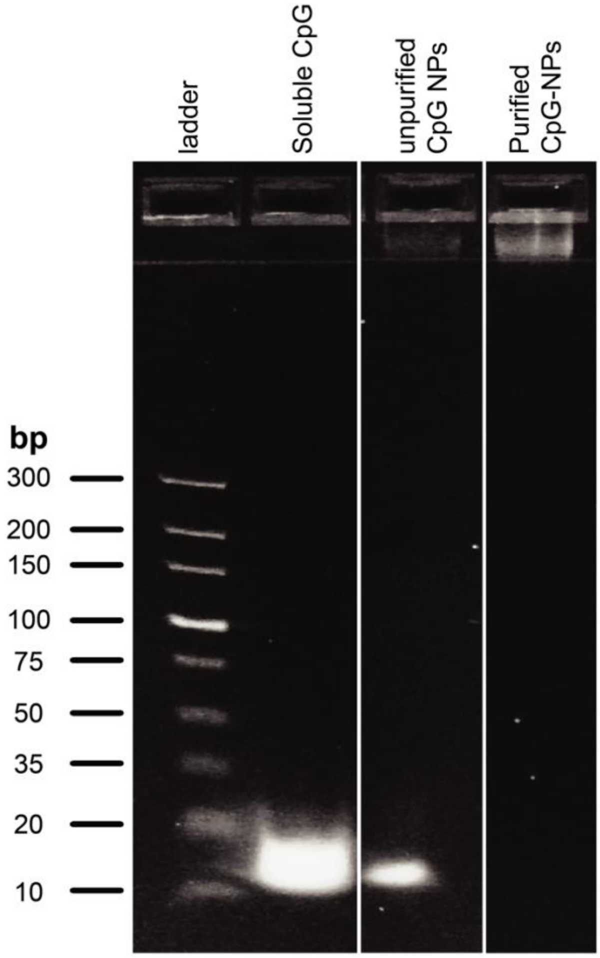


**Figure S2:** Gel electrophoresis of CpG-NPs. Gel electrophoresis after purification of the CpG-NPs demonstrates complete removal of free CpG from the NPs suspension. Soluble CpG runs through the gel and confirms the 20 base pair length. Unpurified CpG-NPs show presence of both, CpG-NPs in the wells (top) and soluble CpG migrating in the gel. Purified CpG-NPs stay in the well of the agarose gel, consistent with NPs conjugation and purification.

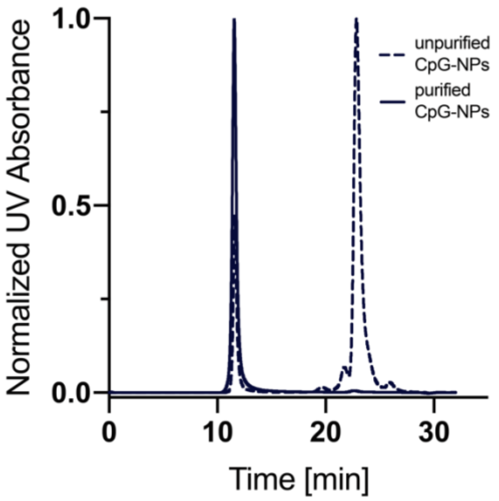


**Figure S3:** GPC traces of 30% CpG-NPs before (3 molar excess of DBCO-CpG) and after purification through a SEC column. Disappearance of the DBCO-CpG peak at 23 mins confirms complete removal of the unconjugated soluble CpG.

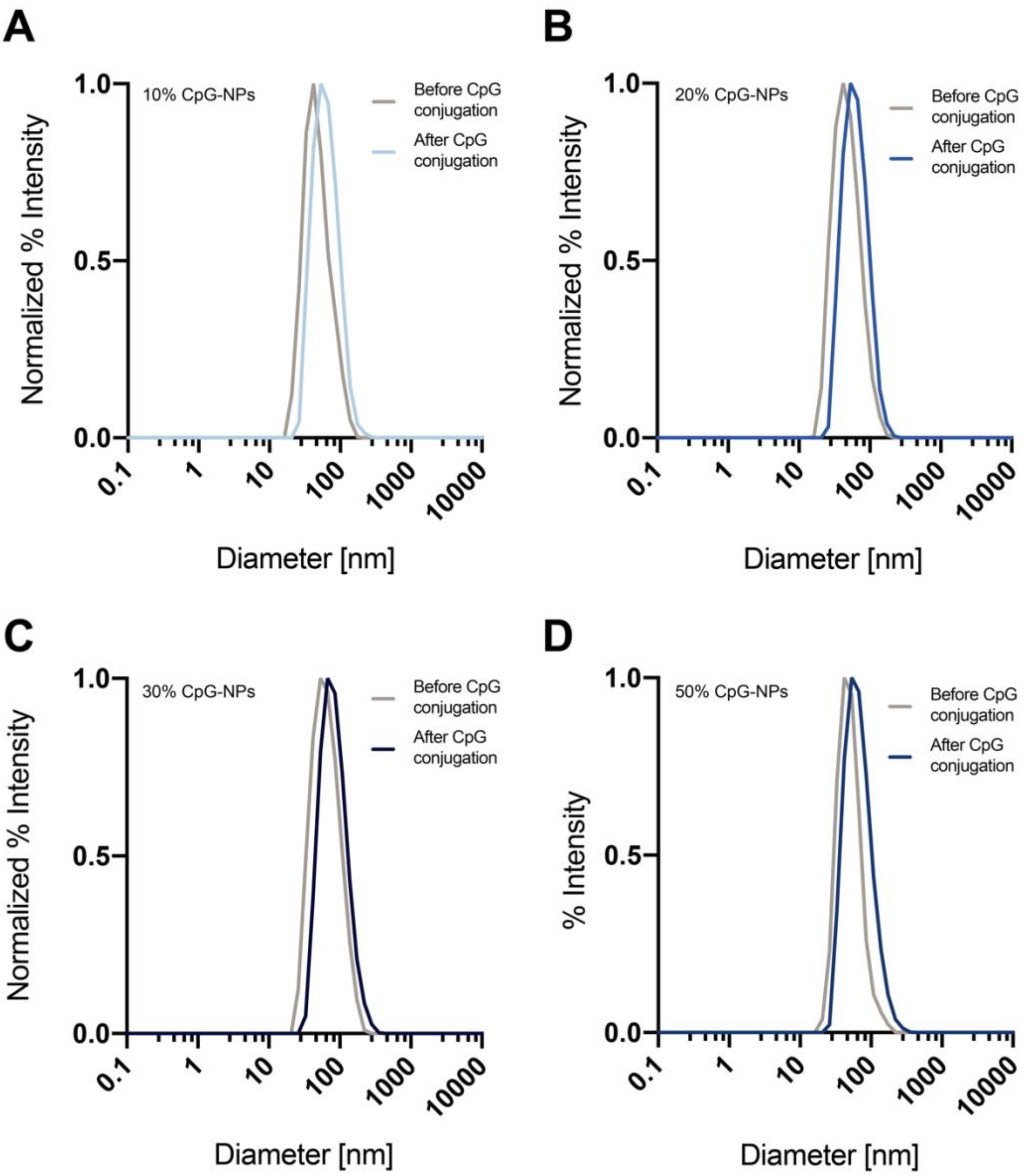


**Figure S4:** Representative dynamic light scattering (DLS) curves of CpG-NPs. Curves show size distribution before (grey) and after CpG conjugation (blue) of (A) 10% valency, (B) 20% valency, (C) 30% valency, (D) 50% valency.

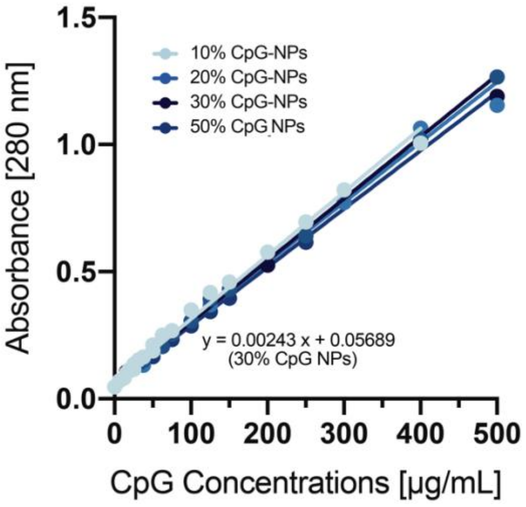


**Figure S5:** Calibration curves of CpG-NPs. CpG-NPs absorbance was measured at λ = 280 nm using soluble CpG. Calibration curves are used to measure the exact concentration of CpG on the NPs with different valencies. CpG concentration is meant the total concentration of CpG conjugated to the NPs in 60 μL of CpG-NPs solution.

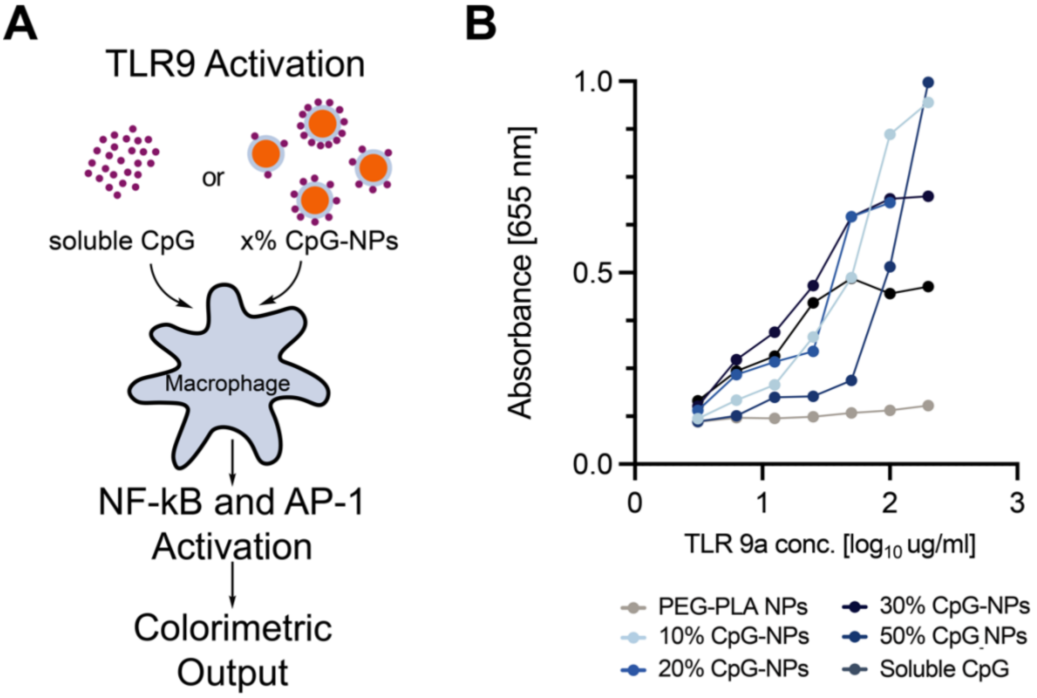


**Figure S6: In vitro activity of CpG functionalized NPs. (A)** Incubation of Raw-Blue macrophage cells (APCs) with either free CpG or different valencies of CpG-NPs (10%, 20%, 30%, 50%) induces the activation of NF-kB and AP-1. The magnitude of activation is quantified via calorimetric output using QUANTI-Blue solution. **(B)** Raw value activation curves across a range of CpG concentrations (3.1–29 µg/mL) delivered on CpG-NPs at different densities to 100,000 Raw-Blue cells. The absorbance at 655 nm corresponds to TLR activation.

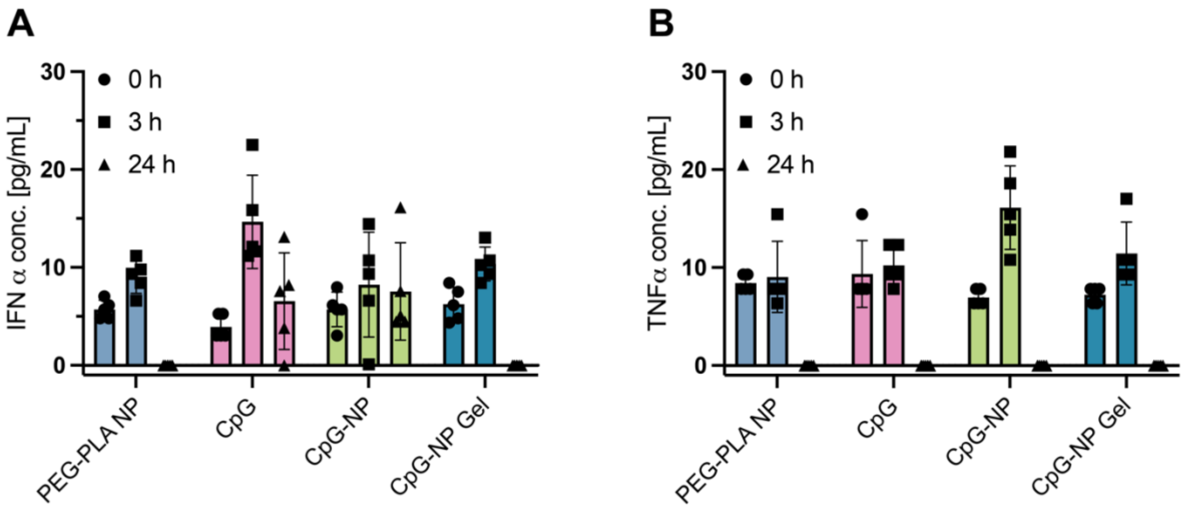


**Figure S7:** Analysis of systemic toxicity. ELISA analysis of (A) IFN-α and (B) TNF-α serum at 0 h, 3 h, and 24 of CpG adjuvanted vaccines with CpG being either in soluble form (CpG), tethered to the NPs (CpG-NP) or tethered to the NPs and encapsulated in the hydrogel (CpG-NP gel) (n= 5).

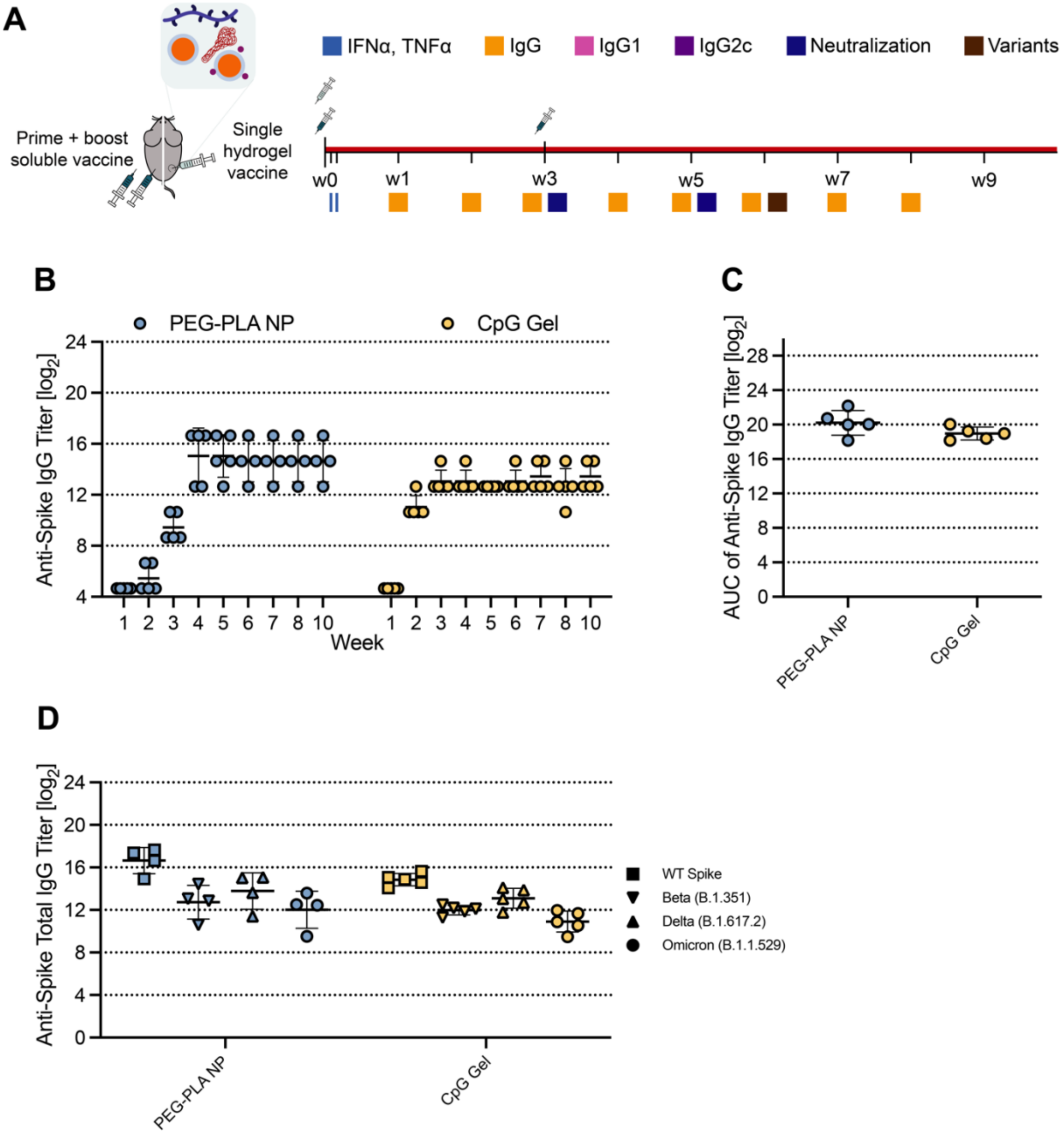


**Figure S8: In vivo humoral response to COVID-19 subunit vaccine. (A)** Timeline of mouse immunizations and blood collection for different assays. Control PEG-PLA NP soluble vaccine group was immunized with a prime dose of 10 µg spike antigen and 20 µg PEG-PLA NP at day 0 and received a booster injection of the same treatment at day 21. CpG Gel group was immunized with a prime dose of 20 µg spike antigen and 40 µg of soluble CpG and did not received a boost. Serum was collected over time to determine cytokine levels and IgG titers. IgG1, IgG2b, and IgG2c titers were quantified and neutralization assays were conducted on day 21 and day 35 serum. **(B)** Anti-spike total IgG ELISA endpoint titer of the vaccine before and after boosting. **(C)** Area under the curve (AUC) of anti-spike titers from (B). **(D)** Anti-spike IgG ELISA titers from serum collected week 6. Titers were determined for wildtype spike as well as Beta (B.1.351), Delta (B.1.617.2, and Omicron (B.1.1.529) variants of spike protein. Each point represents an individual mouse (n = 5). Data are shown as mean +/- s.d.

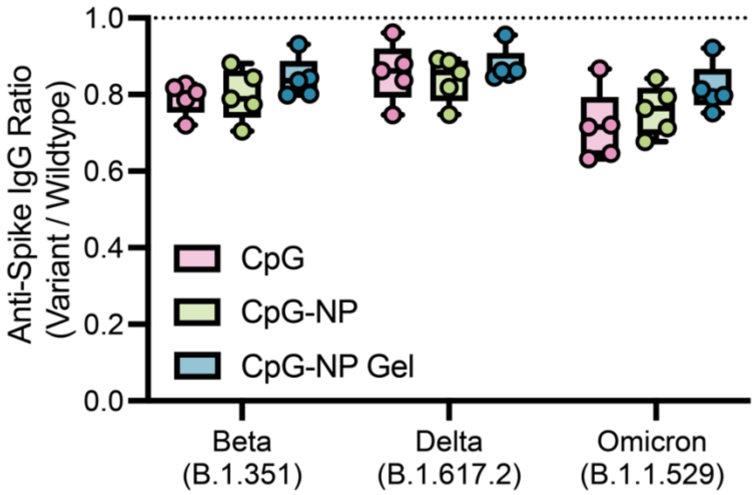


**Figure S9:** Ratio of Anti-Variant Spike IgG titers to Anti-WT spike IgG titers shown in Figure 7D. A ratio closer to 1 indicates a better antibody protection against emerging COVID-19 variants.

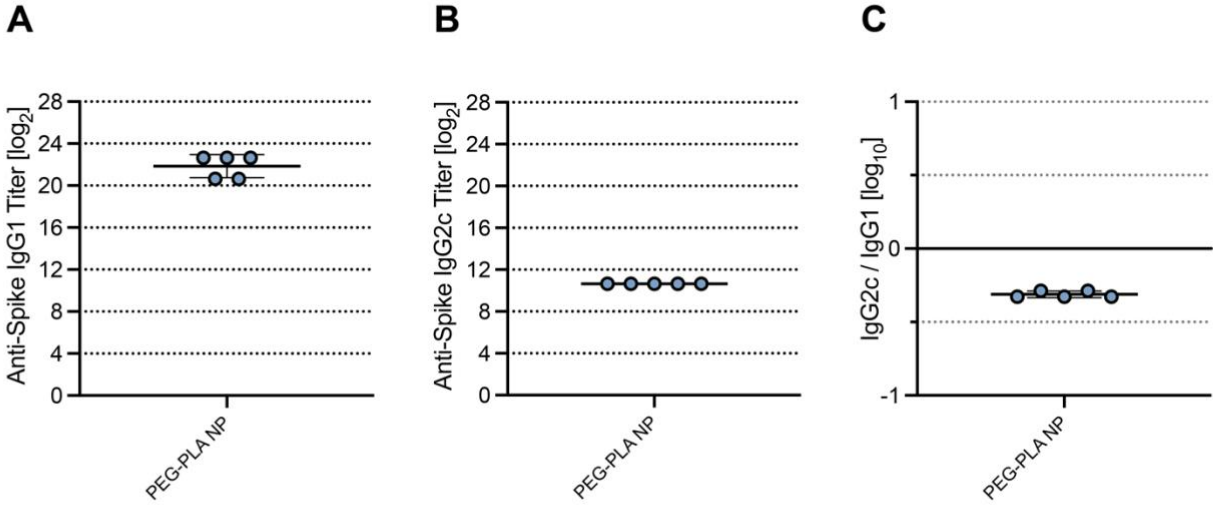


**Figure S10:** Anti-spike IgG1 (**A**) and IgG2c **(B)** titers from serum collected on week 5, 2 weeks after boosting the PEG-PLA NP control group. **(C)** The ratio of Anti-spike IgG2c to IgG1 post-boost titers. Lower values (below 1) suggest a Th2 response or humoral response, and higher values (above 1) suggest a Th1 response or cellular response.

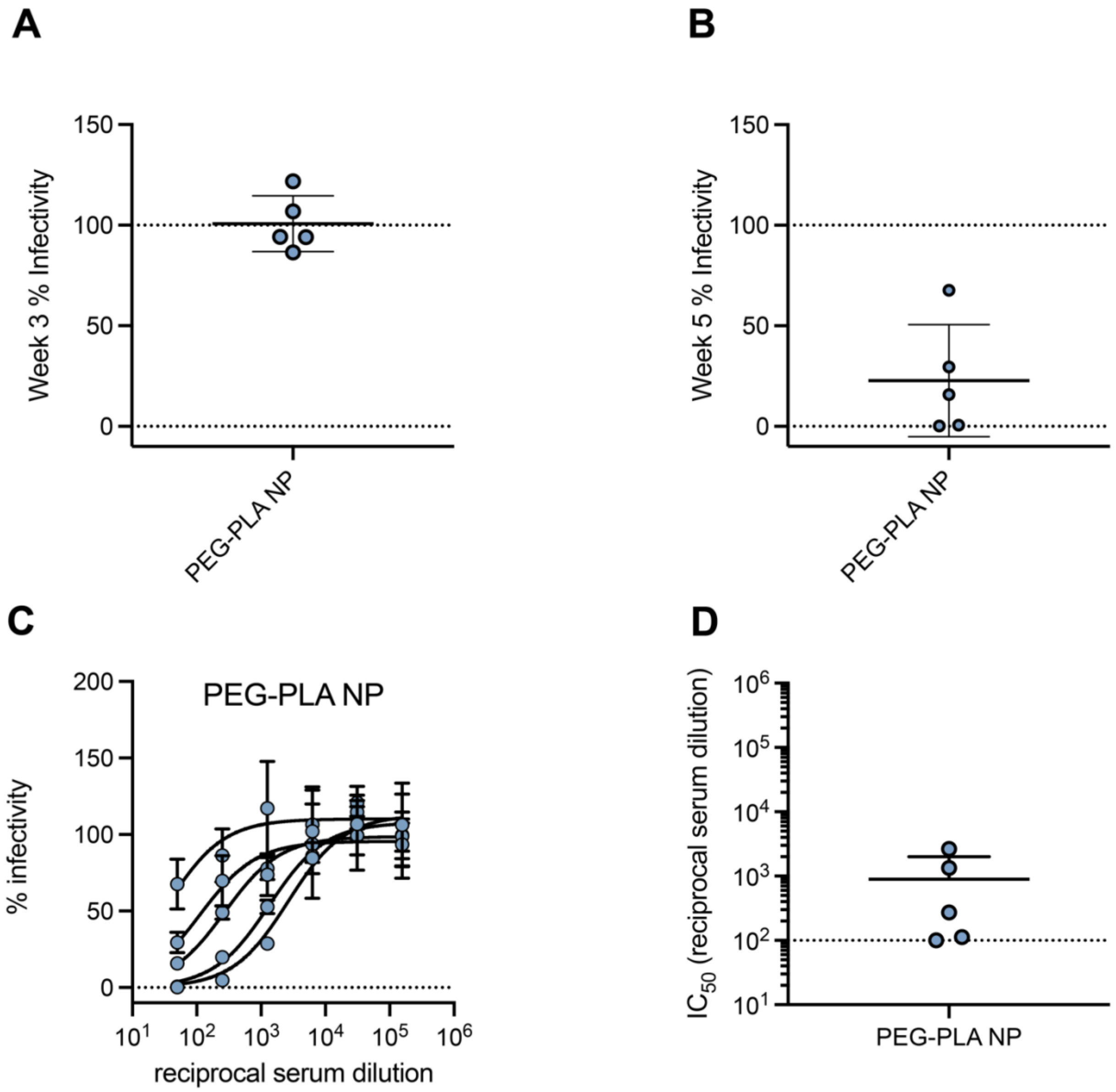


**Figure S11: (A)** Pre-boost (Day 21) spike-pseudotyped viral neutralization assays for the PEG-PLA NP COVID-19 spike vaccine at a serum dilution of 1:50. **(B)** Post-boost (Day 35) spike-pseudotyped viral neutralization assays at a serum dilution of 1:50. **(C)** Percent infectivity at a range of Week 5 serum dilutions as determined by a SARS-CoV-2 spike-pseudotyped viral neutralization assay. **(D)** $IC_{50}$ values determined from neutralization curves on Day 35 following immunization with PEG-PLA NP COVID-19 spike vaccines. Each point represents an individual mouse (n = 5).

**Table S1:** Diameter and PdI of PEG-*b*-PLA and (10%, 20%, 30%, 50%) CpG-NPs

| | Diameter (nm) | PDI |
|---|---|---|
| **PEG-*b*-PLA NPs** | 42.0<br>41.9<br>41.9 | 0.078<br>0.081<br>0.090 |
| **10% CpG-NPs** | 57.5<br>56.0<br>57.1 | 0.219<br>0.220<br>0.225 |
| **20% CpG-NPs** | 59.4<br>60.0<br>59.4 | 0.192<br>0.189<br>0.191 |
| **30% CpG-NPs** | 62.1<br>62.7<br>62.1 | 0.175<br>0.162<br>0.177 |
| **50% CpG-NPs** | 56.6<br>56.7<br>56.1 | 0.207<br>0.222<br>0.178 |
